# Using Machine Learning Enabled Phenotyping To Characterize Nodulation In Three Early Vegetative Stages In Soybean

**DOI:** 10.1101/2022.09.28.509969

**Authors:** Clayton N. Carley, Melinda Zubrod, Somak Dutta, Asheesh K. Singh

## Abstract

The symbiotic relationship between soybean [*Glycine max* L. (Merr.)] roots and bacteria (*Bradyrhizobium japonicum*) lead to the development of nodules, important legume root structures where atmospheric nitrogen (N_2_) is fixed into bio-available ammonia (NH_3_) for plant growth and development. With the recent development of the Soybean Nodule Acquisition Pipeline (SNAP), nodules can more easily be quantified and evaluated for genetic diversity and growth patterns across unique soybean root system architectures. We explored six diverse soybean genotypes across three field year combinations in three early vegetative stages of development and report the unique relationships between soybean nodules in the taproot and non-taproot growth zones of diverse root system architectures of these genotypes. We found unique growth patterns in the nodules of taproots showing genotypic differences in how nodules grew in count, size, and total nodule area per genotype compared to non-taproot nodules. We propose that nodulation should be defined as a function of both nodule count and individual nodule area resulting in a total nodule area per root or growth regions of the root. We also report on the relationships between the nodules and total nitrogen in the seed at maturity, finding a strong correlation between the taproot nodules and final seed nitrogen at maturity. The applications of these findings could lead to an enhanced understanding of the plant-*Bradyrhizobium* relationship, and exploring these relationships could lead to leveraging greater nitrogen use efficiency and nodulation carbon to nitrogen production efficiency across the soybean germplasm.

**Core Ideas:**

- The growth and development of soybean nodules on the taproot and non-taproots have unique growth and development patterns.
- In general, taproot nodules increase in area, while non-taproot nodules increase in count and area.
- Nodulation should be defined by the total nodule area as a function of both nodule count and individual nodule size.
- Genotypes adjust their nodulation through either increasing nodule count or nodule size to increase the total nodule area per root between each growth stage.
- There is a strong correlation between early growth stage taproot nodules and final seed nitrogen content.

## 1. INTRODUCTION

The production of ammonia (NH_3_) through the Haber-Bosch system uses enormous energy to convert atmospheric nitrogen (N_2_) into an available ammonia form as fertilizers, which is required for plant health, yet symbiotic bacteria in legume plants have been doing this for over 60 million years (Mylona et al., 1995; Sprent, 2007). The specialized root structures, known as nodules, are where symbiotic bacteria species fix atmospheric nitrogen into ammonia, which is bioavailable to the plant and essential to meet its growth and developmental needs, including metabolic needs, protein, and amino acids production. The seeds of nitrogen-fixing legume crops typically have 14-36% crude protein compared to major cereal grain crops with 6-11% (Akibode & Maredia, 2012; Hall et al., 2017; Koehler & Wieser, 2013). In crop species that can form nodules, N fixation allows the host plant to meet up to 98% of its nitrogen requirements depending on the prevailing soil type (Bergersen et al., 1985; Hardarson et al., 1984; Matheny & Hunt, 1983; Roy et al., 2020; Salvagiotti et al., 2008).

There are several ways to determine N-fixation rates. In soybean, the more common methods are through the study of nitrate reductase activity (Fabre & Planchon, 2000; Soussana et al., 1989; Thibodeau & Jaworski, 1975) and ureide measurement (Kibido et al., 2020; Moro Rosso et al., 2021; Takahashi et al., 1992). These methods are instrumental; however, few studies have correlated these chemical N-fixation rates with nodule counts (David F. Herridge et al., 1990). Although nodule count and nodule distribution at single growth stages in numerous environments have been studied (Caetano-Anolles & Gresshoff, 1993; Chung et al., 2020; Kluson et al., 1986; Voorhees et al., 1976), relatively few studies have evaluated nodulation across multiple growth stages (Fabre & Planchon, 2000; David F. Herridge et al., 1990). A knowledge gap exists between N-fixation with nodule counts and their placement throughout the roots in the early vegetative stages (Fabre & Planchon, 2000). Prior research has established the critical role of nodulation in early growth stages on chlorophyll content, leaf size, plant height, number of pods per plant, 1000-seed weight, and seed protein and oil content (Vollmann et al., 2011); therefore, it is imperative to explore nodule formation and distribution on roots at early growth stages.

While the process of nodulation and N-fixing has garnered scientific interest and research (M. Du et al., 2020; Ren et al., 2019), there is still a gap in the development of methods to quantify nodules in legume crops. Limited attempts have been made to quantify nodule size and numbers through field environments due to the exorbitant requirements for labor and work to phenotype nodules in a reasonable scale and time. Recent and continuous advancements in deep learning and computer vision for plant features and organ detection (Akintayo et al., 2018; Falk, Jubery, Mirnezami, et al., 2020; A. Singh et al., 2016, 2021; Asheesh Kumar Singh et al., 2018) enable further work in phenomics to ask more detailed and unique questions (Bucksch et al., 2014; Falk, Jubery, Mirnezami, et al., 2020; Riera et al., 2021; Seethepalli et al., 2020; Asheesh K. Singh et al., 2021; Smith et al., 2020). The use of machine learning also makes trait acquisition more consistent and feasible (Jiang & Li, 2020; Riera et al., 2021) for complex traits such as disease detection (Ghosal et al., 2018; Nagasubramanian et al., 2018), small object detection for soybean cyst nematode (SNC) eggs (Akintayo et al., 2018), and abiotic stress (Naik et al., 2017; J. Zhang et al., 2017). Advances in image- and ML-based methods provide exciting solutions for root trait and nodule related studies.

The number of nodules can vary on soybean roots from a few to several hundred per plant, depending on the genotype, growth stage, and root system architecture (Harris & Dickstein, 2010; Huault et al., 2014; Yang et al., 2017). Yet, super nodulating plants are not the most efficient in nitrogen use (Kohl et al., 1980; Weber, 1966; Wu & Harper, 1991). Manual measurements and counts are inefficient due to the complexity of the number, size, and placement of nodules on roots. Therefore, automated methods, including computer vision and machine learning-based models, are valuable avenues to solve these challenges (Kar et al., 2022; Mochida et al., 2019; Nagasubramanian et al., 2022; Asheesh K. Singh et al., 2021). While some computer vision techniques have been used to identify nodule patterns in semi-controlled environments (Barbedo, 2012; S. Han et al., 2015; Remmler et al., 2014), only recently have attempts been made to accurately field phenotype and quantify nodulation responses using machine learning (Chung et al., 2020; Jubery et al., 2021). The development of the Soybean Nodule Acquisition Pipeline (SNAP) (Jubery et al., 2021) has enabled the accurate quantification of nodule location and size on roots at the early vegetative growth stages. Exploring the relationships between nodule development, size, and root location will be useful for breeding approaches and can further explore the balance and contribution of nodulation to nitrogen use efficiency across diverse soybean lines.

There have been numerous links between genetic controls for root system architecture (RSA) and nodulation. Within *Medicago truncatula*, a model plant for legume species, there is evidence that nodulation rates can be linked with RSA as the *ACA2* gene negatively regulates lateral root formation while positively controlling symbiotic nodulation (Huault et al., 2014), and the *LATD*/*NIP* gene is essential for nodule meristems as well as the development of lateral and primary roots (Harris & Dickstein, 2010). Further work in Soybeans has shown that a micro-RNA, *miR2111*, is a component of root-to-shoot signaling that positively regulates root nodule development and shapes lateral root emergence RSA (M. Zhang et al., 2021). While work has been done to evaluate the changes and controls of RSA traits and their impacts on overall nodulation, little has been done to assess the differences in nodulation across the component parts of RSA, such as nodulation distinctions between the primary taproot and lateral secondary non-taproots.

There are several reasons that a time series investigation of nodules, their size, and placement is needed. Since nodules are dynamic organs growing on roots that both develop and senesce, there are numerous hormones, redox signals, and environmental triggers that can control or impact their growth and development (Kazmierczak et al., 2020; Marquez-Garcia et al., 2015; Puppo et al., 2005). The lifecycle and development of these nodules are also dependent on the type of bacterium that develop within the nodule, which can be dependent on local bacterium availability, soil type, and soil pH (Q. Han et al., 2020). The nitrogen fixed through soybean nodulation has been shown to have a vast range based on available nitrates, mineralized N, weather, and soil type and can range from 0 to 337 kg ha^-1,^ which has been shown to be up to 98% of the total nitrogen demands of the plant (Ciampitti & Salvagiotti, 2018; Córdova et al., 2019; D. F. Herridge, 1982; Weber, 1966). The fixation of this nitrogen has dramatic energy needs as 33% of the root translocated photosynthate carbon can be utilized by early developing nodules alone and up to 50% during optimal N fixation in later growth stages (Warembourg et al., 1982). Being able to adapt and optimize the nodulation relationships between carbon consumed and nitrogen fixed could dramatically impact the yield-protein quality breeding strategies of soybeans as it has been calculated with the Kjeldahl conversation method that seed protein is 5.66-5.79 times the total N concentration of mature seed (Morr, 1982), which on average requires 2-2.25 kg N per 27.2 kg of soybeans (4.5 - 5 lbs. of N bu^-1^) to produce. As extensive amounts of nitrogen are required for vegetative growth and seed fill (Basal & Szabó, 2020), exploring the relationship between nodulation and nitrogen content within the plant enables a deeper understanding of soybean nitrogen use efficiency (Hao et al., 2011).

With these motivations, we performed temporal field evaluations of six diverse soybean lines chosen for their root system architecture differences (Falk, Jubery, O’Rourke, et al., 2020; Falk, 2019). The objectives of this study are to: (1) Explore and identify the trends and relationships between soybean nodules and early growth stage developments, (2) Quantify the nodule counts, individual nodule area, and total nodule area between two unique spatial locations of the taproot and non-taproot growth zones, and (3) Identify relationships between the traits mentioned above and the amount of nitrogen in the seeds of mature plants. We report the variability of nodule development in these early growth stages between the non-taproots and taproots and the nodule relationship with above and below-ground biomass and seed nitrogen content. Our results indicate that while non-taproot root nodules increase in count between each early growth stage for all tested genotypes, overall, the taproot nodules do not significantly increase in count after the V1 growth stage. Further, we found that taproot nodules increase in volume between each growth stage and that some genotypes increase nodule area at greater rates than others. We also report a significant positive correlation specifically between taproot nodules and total nitrogen content in mature seed.

## 2. MATERIALS AND METHODS

### 2.1 Plant Materials

The data collected for nodule growth stage analysis was generated from growing six unique soybean genotypes: CL0J095-4-6, PI 80831, PI 437462A, PI 438103, PI 438133B, PI 471899 (USDA, 2022). The six lines evaluated in this study were selected due to their unique phenotypic characteristics. They were chosen to represent a broad diversity of country of origin, genotype, and historic yield phenotypes. These genotypes were selected from the USDA mini core collection based on similar developmental time stages using historical data to align early vegetative growth stages, flowering, and physiological maturity. From the lines with similar developmental stages, an identical-by-state Van Raden kinship matrix (VanRaden, 2008) was calculated to select lines that were genetically dissimilar to each other. Seven genetically similar clusters were observed among the soybean collection, and visual observations were made of the root phenotypes in each cluster to identify the most phenotypically diverse lines for selection (Falk, Jubery, O’Rourke, et al., 2020). CL0J095-4-6 was found to have a considerably robust quantity of lateral roots and a consistently large taproot with average nodulation. PI 437462A had large lateral roots near the top of the taproot but fewer lateral roots lower in the profile with low nodule counts. PI 438103 had an average root mass but with large nodules. PI 438133B had an average number of lateral roots, but they were consistently long and had higher visual nodule counts. PI 471899 had few lateral roots and few nodules with more nodules on the taproot. PI 800831 had large nodules and very steep root angles from the tap root. The yield phenotypes used were the observed means from data collected in 2017 across four environments in Iowa (Parmley et al., 2019). PI 437462A originating from Russia, had a relatively low calculated yield at 1143 kg/ha (17 Bu/A), while three lines, PI 438103, PI 438133B, and PI 80831, originate from China and represent an intermediary calculated seed yield of 1749, 2286, 1749 kg/ha (26, 34, and 26 Bu/A) respectively. PI 471899 was selected as a high yielding PI line from Indonesia with a calculated yield of 3766 kg/ha (56 Bu/A), and CL0J095-4-6 was chosen as a high yielding line from the United States with a calculated yield of 3160 kg/ha (47 Bu/A). CL0J095-4-6 is also one of the parental strains of the soybean nested association mapping set (Diers et al., 2018). Visual representation of the selected roots can be found in supplemental figure S1.

### 2.2 Root Excavation and Experimental Design

This experiment was conducted in three environments (In 2018 and 2019 - Muscatine Island Research Station, Fruitland, IA; Soil type - Fruitland Coarse Sand. In 2018 - Horticulture Research Station, Gilbert, IA; soil type: Clarion Loam). For ease, we refer to 2018 Horticulture farm as S1, 2018 Muscatine as S2, and 2019 Muscatine as S3. At three early growth stages, V1, V3, and V5, images were collected of each plant root. The SNAP pipeline is currently developed for 2D image and root analysis as the roots typically become too rigid by R1. The design at each environment is a split-plot experiment with a whole-plot factor (growth stages nested within environments) and a split-plot factor (genotypes). The design at the whole plot level is completely randomized, and the design at the split-plot level is a randomized complete block design with ten to twelve replications of each block. There are two types of experimental units in a split-plot experiment: whole-plots and split-plots. The split-plots are “plots” (with a single plant per plot) planted 100 cm × 100 cm apart. Three seeds per plot were initially planted, but after emergence, two were cut using a sharp blade to leave one standing plant per plot. At the appropriate growth stage, plants were tagged with a barcode and labeled with identification strips. The roots were extracted using trenching spades (120 cm steel round drain spade, wood handle) from a 50-cm diameter and 30-cm deep area. The plant roots were dug with extreme caution not to break roots or dislodge the nodules during extraction. This was followed by gently removing the roots from the soil by hand, ensuring little to no damage to roots and nodules. When initially evaluating the removal methods, a thorough search of the soil after root removal was conducted and found insignificant nodule loss (less than 5% of total nodules on the roots regardless of size), suggesting that any incidental loss incurred would have minimal influences on the study. Furthermore, none of the six genotypes showed any proclivity for losing more or fewer nodules than the other.

### 2.3 Imaging Protocols

Details on imaging protocols are provided in (Jubery et al., 2021). In brief, five-gallon buckets half full of water were used to transport and rinse the remaining soil from the roots after extraction (See section 2.2). Next, the roots were placed on trays that had been painted blue for a consistent background measuring 35 × 50cm with a 2 cm lip. The imaging trays were filled halfway with water, and the roots were gently separated to prevent increased occlusion or clumping of the roots in each image. To obtain a precise 2D image of each plant root, a glass plate fitted to the size of the tray was then placed on top of the root in the water to hold it in place, and just enough water to cover the plate was added. The plate was then slid into an imaging platform customized for this process.

Aluminum T-slot extrusion frames (80/20 Inc., Columbia City, IN) were used to build the platform with two softbox photography lights (Neewer; Shenzhen, China). A Canon T5i digital SLR camera (lens: EF-S 18-55mm f/3.5-5.6 IS II) (Canon USA, Inc., Melville, NY) was mounted 55cm above the imaging plane, and four 70-watt CFL bulbs were used to provide consistent illumination. It was found that placing a black sheet over the top of the entire platform prevented undesired room lights from reflecting into the images. It was tethered to a laptop operating Smart Shooter 4 (Hart, 2019) photo capturing and camera control software to trigger capture and automatically extract the barcodes placed at the top of each image from the plant tags for image labeling and naming. For additional details, see (Falk, Jubery, O’Rourke, et al., 2020).

Following root imaging, the dry mass of the roots and shoots was acquired by drying the roots in paper bags at 60 degrees C for two days. The shoots were placed in separate bags for concurrent root shoot drying. After the roots and shoots were dehydrated, the roots were weighed for dry mass, including nodules. Nodules from each root were removed by trained researchers and counted for ground truth validation of the SNAP pipeline. The total nodule weight per root was measured in grams along with the total shoot biomass weight.

### 2.4 Image Analysis with SNAP and Data Cleaning

Each image was run through the SNAP as described and explained in Jubery et al. to extract the quantified nodule counts, individual nodule sizes, and locations on each root (Jubery et al., 2021). The extracted data was then visually assessed for any taproot or nodule misclassifications. Any taproots that were misidentified were hand-annotated, and a manual binary mask of the taproot was created for these roots to generate accurate taproot nodule metrics. Additionally, any nodules calculated as greater than 1 cm^2^ were visually assessed for misclassification and removed from the data set if they were false positives. This resulted in two out of 50,848 nodules being removed. Any nodules less than .001 cm^2^ were removed from the dataset as false positives. Under this criterion, four nodules out of 50,848 nodules were removed. By Subtracting the taproot nodule data from the total nodule data, we imputed the non-taproot nodule information. Important traits that were collected through SNAP for the taproot and non-taproot growth zones include (a) individual nodule count, which is the physical count of nodules from SNAP, (b) individual nodule area, which is the average nodule size of each nodule on a given root, and (c) total nodule area which is the sum of all nodule areas on a given root.

### 2.5 Measuring Percent N in Mature Seed

Seed was collected from mature plants (R8) in three replications from each environment to gather data on the amount of nitrogen in the mature seed. The seed from each plant was bulked and ground into a fine powder. This was done by weighing 5 grams of soybeans to the nearest whole bean over 5 grams. Then the seeds were added to a Mr. Coffee IDS 57 grinder, grinding them for 30 seconds, then pulse grinding the samples for 30 seconds while tapping the grinder walls to ensure that the sample was fully homogenized. A spoonula was then used to remove the milled bean material from the grinder and was placed into a 15 mL sterile centrifuge tube. Distilled water was then used to triple rinse the grinder, the grinder was pulsed with water inside to ensure blade cleanliness, and excess water was wiped from the grinder with a clean paper towel. The grinder was then rinsed with 80% V/V ethyl alcohol, and excess alcohol was wiped out of the grinder and air-dried before the next sample. The spoonula was triple rinsed with distilled water, sprayed with alcohol, and excess alcohol was wiped off. A thorough visual examination was conducted to ensure no excess soybean material was left in the grinder or on the spoonula. The samples were sent for elemental analysis to obtain data on the amount of elemental nitrogen, carbon, hydrogen, and sulfur in each sample. Analysis was performed using a Thermo FlashSmart 2000 CHNS/O Combustion Elemental Analyzer in the Chemical Instrumentation Facility at Iowa State University.

### 2.6 Statistical Analysis

Analysis was conducted using R statistical software (Ripley, 2001). A mixed linear model was used for analysis and fit using the R package lme4 (Bates et al., 2015) with further analysis using emmeans (Lenth, 2022). Upon initial review of the raw data, heteroscedasticity was observed in the residual means of the nodule count and taproot nodules evaluated. Therefore, a log transformation was conducted and chosen to evaluate the data moving forward. Furthermore, due to instances (less than 5 out of 542, prior to outlier removal) of taproot and non-taproot nodule counts being zero, the log transformation of dependent variables (i.e., taproot nodules and non-taproot nodules) was conducted as log (Nodule Count + 1) to ensure all data were included in the analysis. For each explored trait, a mixed linear model was fit with the form using the log transformed data:

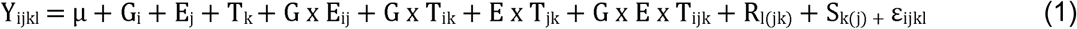

Where *y* is a vector of observed phenotypes, μ is the grand mean, *G*_*i*_ is the effect of the *i*^*th*^ of six genotypes, *E*_*j*_ is the effect of the *j*^*th*^ of three environments, and *T*_*k*_ is the effect of the *k*^*th*^ of three growth stages. *G x E*_*ij*_ is the interaction between the *i*^*th*^ genotype and the *j*^*th*^ environment. *G x T*_*ik*_ is the interaction between the *i*^*th*^ genotype and *k*^th^ growth stage. *E x T*_*jk*_ is the interaction between the *j*^*th*^ environment and the *k*^*th*^ growth stage. *G x E x T*_*ijk*_ is the interaction between the *i*^*th*^ genotype, *j*^*th*^ environment, and the *k*^*th*^ growth stage. *R*_*l(jk)*_ is the random effect of the *l*^*th*^ replicate nested within the *j*^*th*^ environment, and *k*^*th*^ growth stage, *S*_*k(j)*_ is the random effect of the *k*^*th*^ growth stage nested within the *j*^*th*^ environment, and ε*ijkl* is the residual error, and is assumed to be normally and independently distributed, with mean zero and variance ^2^. The observed phenotypes of nodule counts, including taproot and non-taproot counts along with root and shoot biomass, were log-transformed for analysis using the natural log of *y*_*ijkl*_ in model 1.

Outliers were selected and removed after calculating the studentized residuals using the ‘stats::rstudent’ function(Fox & Weisberg, 2011), where outliers were excluded when they had a standardized residual being greater than 3.23 as calculated in Lund (1975)(Lund, 1975) with α = This resulted in 9 of the original 542 roots being identified as outliers, including four at V1, three at V3, and two at V5 roots. Of these nine roots, after an in-depth manual observation, the outlier roots were found to have architecture or image anomalies and provided further evidence for the decision to remove these nine roots for all analyses. Any outliers identified for the remaining biological biomass traits, i.e., root and shoot biomass, were excluded from further analyses. The following sample quantities were excluded in each dataset category due to collection errors or outlier status - root dry mass, 22 out of 542; shoot dry mass, 11 out of 542.

To conduct a type III analysis of variance in R, we used lme4 (Bates et al., 2015) and lmerTest (Kuznetsova et al., 2017) and calculated the denominator degrees of freedom for F statistics with the Kenward and Roger method (Kenward & Roger, 1997). Correlations were assessed using Pearson’s correlations of the mean values at each environment and growth stage for each trait against the other using the R “Hmisc” package and “rcorr” function (Harrell, 2021). For comparing traits against the final seed nitrogen content, correlations with significance greater than α = 0.1 were reported.

## 3. RESULTS

### 3.1 Diverse genotypes express variable root phenotypes across environments and years

The analysis of variance of non-taproot nodule counts and taproot nodule counts is presented in Table 1. Genotype, environment, growth stage, and genotype-by-environment interactions were significant in the non-taproot nodules. Additionally, the genotype by growth stage and environment by growth stage interactions were significant, showing that the genotype and environment both have unique interactions with the growth stage. Only the genotype and genotype by environment interactions were significant for taproot nodules. Similar to the non-taproot nodules, these genotypes have varying rates of nodule counts between them and showed significant interactions with the environments they are grown in. The magnitude of the effect of environment on taproot nodules was less than non-taproot nodules suggesting that the taproot nodules are more stable across environments compared to genotypes.

**Table 1.** Analysis of Variance of Non-Taproot and Taproot nodules with Kenward-Roger’s method.

| Source of variation | dF | Taproot Nodules |  | Non-Taproot Nodules |  |
| --- | --- | --- | --- | --- | --- |
|  |  | F Value | P value | F Value | P Value |
| Genotype (G) | 5 | 33.81 | $<2.2 \times 10^{-16}***$ | 26.94 | $<2.2 \times 10^{-16}***$ |
| Environment (E) | 2 | 2.25 | 0.10* | 12.85 | $2.67 \times 10^{-6}***$ |
| Growth Stage (T) | 2 | 0.29 | 0.74 | 19.75 | $2.72 \times 10^{-9}***$ |
| G x E | 10 | 7.49 | $5.72 \times 10^{-11}***$ | 4.59 | $3.45 \times 10^{-6}***$ |
| G x T | 10 | 1.52 | 0.13 | 1.62 | 0.09* |
| E x T | 4 | 0.19 | 0.95 | 2.38 | 0.04** |
| G x E x T | 20 | 1.21 | 0.24 | 1.66 | 0.03** |

Significance at \*\*\* < 0.001, \*\* 0.05, \*0.1

We observed differences in the nodule counts, nodule taproot counts, and root biomass at each growth stage across the three environments (Figure 1). The traits from S2 were numerically lower than S1 and S3, aside from the V3 root biomass. We observed similar trends in the V5 stage (Figures 1A and 1C), but compared to V5, a substantial increase in overall nodules and biomass in V5 at S1 was noted (Figure 1B). In contrast, the taproot nodules were consistent with the other two environments. It was also observed that, at S2, all three traits were lagging but caught up by V5. We also noted that while biomass is consistently increasing, a lower unproportional increase is observed in total nodules and taproot nodules. We also observe increases between growth stages in total nodules but not in taproot nodules.

**Figure 1.**
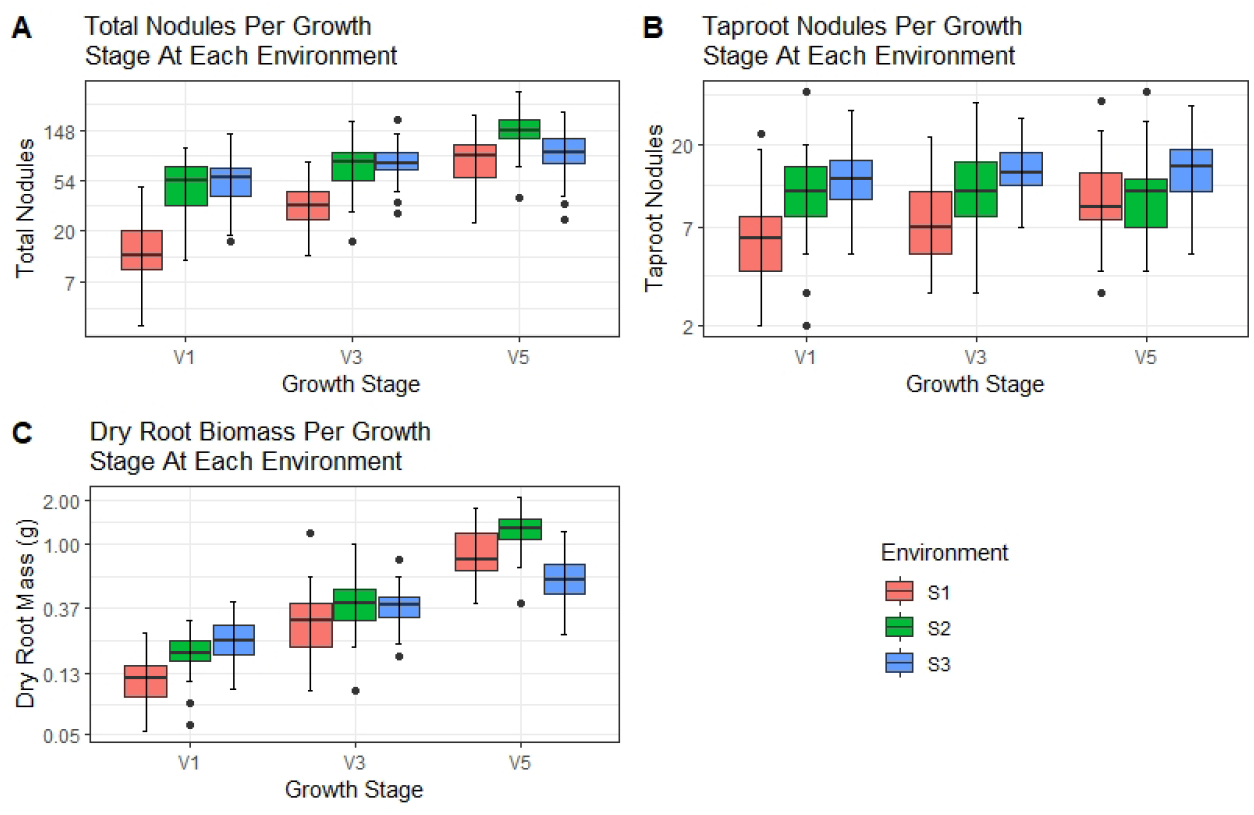
Each environment shows variability in the overall means of total nodule counts and biomass between the environments for each trait in soybean, with data compiled from six diverse genotypes. a) Boxplots showing the overall total nodules per root means of all genotypes at each growth stage and environment on a y-axis log scale. b) Boxplots showing the overall means of the nodules on the taproots in all genotypes at each growth stage and environment on a y-axis log scale. c) Boxplots showing the overall means of the dried root mass in grams from all genotypes at each growth stage and environment on a y-axis log scale.

In tables 2-5, we report the mean and standard error of the collected nodule, biomass, and seed elemental content traits, for each accession and across growth stages at each of the three environments. Table 2 shows the mean count of total nodules and taproot nodules. PI 438133B consistently had the highest mean number of total and taproot nodules. In contrast, PI 471899 generally had the lowest total nodule counts, along with PI 438103, the lowest taproot nodule counts. Table 3 presents each genotype’s total nodule area for taproot and non-taproots in each environment. PI 438133B had a substantially larger total, taproot, and non-taproot nodule area across all three environments. PI 471889 and PI 437426A have the lowest total and non-taproot total nodule areas, and PI 471899 has some of the lowest taproot total nodule areas. PI 437462A has an intermediate taproot total nodule area compared to the other genotypes. Table 4 shows the mean and standard error of the root and shoot dry mass at each growth stage and environment for each genotype. PI 438133B has the highest root and shoot biomasses, while PI 437462A has the smallest. Table 5 shows the mean seed nitrogen, carbon, and one hundred seed weight (Cwt) in mature seed taken at the R8 growth stage for comparison with the other traits. We observed that N and C tend to be more consistent within genotypes while Cwt had more variation between environments.

**Table 2.**
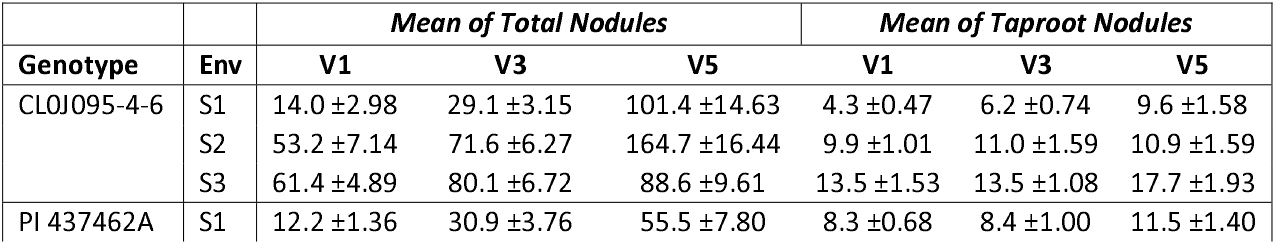
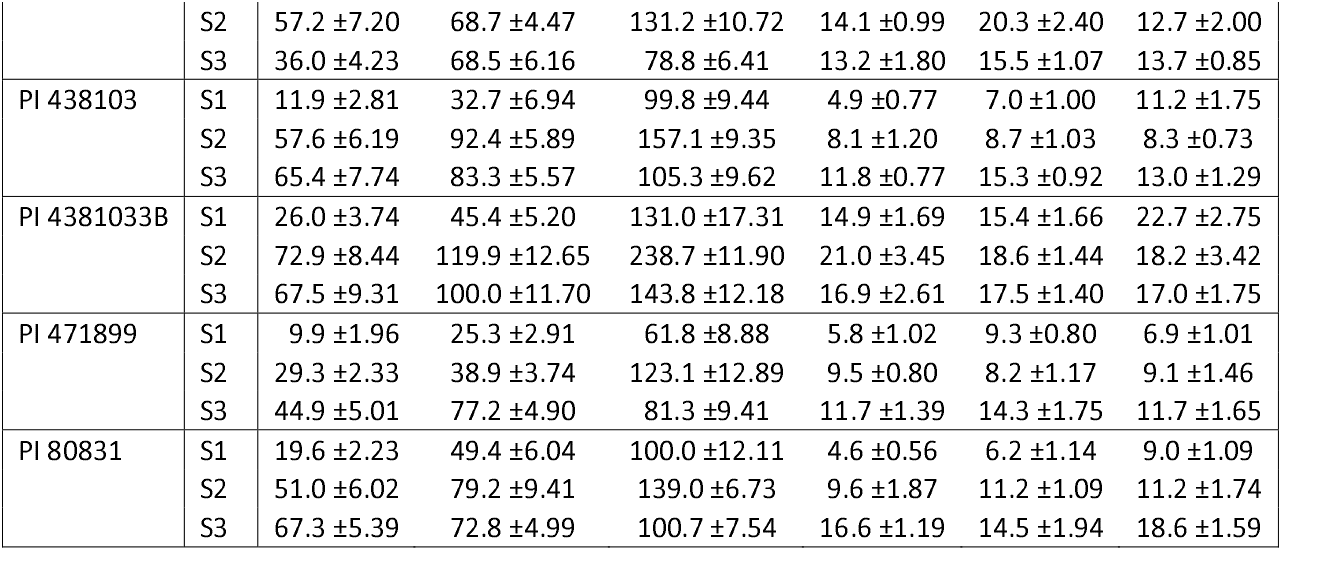
Shows the mean and standard error of the total nodule counts and taproot nodule counts in three vegetative stages for each genotype and across environments.

**Table 3.** Shows the mean and standard error of the total nodule area, taproot nodule area, and non-taproot area per root in three vegetative stages for each genotype and across Three environments.

| Genotype | Env | Mean Total Nodule Area per Root (mm<sup>2</sup>) |  |  | Mean Total Taproot Nodule Area per Root (mm<sup>2</sup>) |  |  | Mean of Total Non-Taproot Nodule Area per Root (mm<sup>2</sup>) |  |  |
| --- | --- | --- | --- | --- | --- | --- | --- | --- | --- | --- |
|  |  | V1 | V3 | V5 | V1 | V3 | V5 | V1 | V3 | V5 |
| CLOJ095-4-6 | S1 | 28.52 ±6.415 | 76.81 ±6.536 | 353.27 ±51.984 | 9.08 ±1.473 | 26.59 ±5.484 | 55.27 ±8.626 | 18.94 ±6.450 | 50.22 ±7.474 | 298.01 ±54.045 |
|  | S2 | 139.59 ±19.633 | 305.51 ±31.842 | 874.82 ±82.731 | 37.22 ±6.493 | 73.37 ±15.951 | 86.41 ±14.419 | 102.37 ±19.120 | 232.13 ±28.362 | 788.42 ±78.970 |
|  | S3 | 187.34 ±14.567 | 317.07 ±27.119 | 445.62 ±52.430 | 74.80 ±10.812 | 83.78 ±8.287 | 122.75 ±9.786 | 112.54 ±13.147 | 233.29 ±24.835 | 322.87 ±53.376 |
| PI 437462A | S1 | 27.78 ±2.965 | 96.33 ±16.185 | 244.55 ±37.665 | 19.48 ±1.621 | 44.07 ±8.04 | 74.80 ±7.747 | 10.37 ±3.002 | 52.27 ±10.822 | 169.75 ±38.839 |
|  | S2 | 139.12 ±15.780 | 265.12 ±20.976 | 684.04 ±70.692 | 51.06 ±4.864 | 102.61 ±11.96 | 101.93 ±17.796 | 88.06 ±14.956 | 162.51 ±23.681 | 582.12 ±66.894 |
|  | S3 | 110.95 ±13.034 | 279.37 ±23.615 | 379.99 ±29.169 | 58.71 ±8.098 | 91.71 ±7.261 | 95.89 ±9.114 | 52.23 ±10.160 | 187.66 ±22.136 | 284.09 ±25.617 |
| PI 438103 | S1 | 18.34 ±4.616 | 68.98 ±13.257 | 380.67 ±45.438 | 8.43 ±1.339 | 21.71 ±3.728 | 57.07 ±9.09 | 11.56 ±5.574 | 47.27 ±14.171 | 323.59 ±38.278 |
|  | S2 | 149.70 ±18.311 | 330.08 ±37.796 | 951.02 ±56.860 | 30.70 ±3.838 | 42.84 ±5.633 | 75.41 ±11.099 | 121.65 ±15.951 | 287.24 ±37.006 | 875.62 ±47.948 |
|  | S3 | 205.28 ±20.845 | 341.22 ±19.078 | 549.45 ±51.786 | 64.35 ±7.343 | 106.83 ±9.417 | 103.56 ±9.74 | 140.93 ±22.244 | 234.40 ±19.979 | 445.89 ±48.345 |
| PI 4381033B | S1 | 66.22 ±14.943 | 175.40 ±21.457 | 696.46 ±107.018 | 41.63 ±4.881 | 79.39 ±9.849 | 165.44 ±24.314 | 9.85 ±2.055 | 96.01 ±19.822 | 531.02 ±112.928 |
|  | S2 | 214.70 ±30.585 | 561.39 ±50.151 | 1381.61 ±83.005 | 80.32 ±8.454 | 124.22 ±9.805 | 149.53 ±24.141 | 134.38 ±31.756 | 437.17 ±53.799 | 1232.09 ±89.723 |
|  | S3 | 262.82 ±25.665 | 416.28 ±44.586 | 739.79 ±70.971 | 99.51 ±15.865 | 117.97 ±9.709 | 111.98 ±10.939 | 145.18 ±23.687 | 298.30 ±38.210 | 627.82 ±67.985 |
| PI 471899 | S1 | 22.31 ±4.190 | 95.52 ±11.834 | 231.17 ±38.105 | 15.43 ±3.214 | 53.24 ±6.171 | 37.58 ±5.960 | 7.73 ±2.706 | 42.27 ±8.726 | 193.59 ±38.161 |
|  | S2 | 95.63 ±6.145 | 172.99 ±17.314 | 614.25 ±54.847 | 42.61 ±4.594 | 50.84 ±9.518 | 62.25 ±12.925 | 53.02 ±7.309 | 122.15 ±22.733 | 606.18 ±30.001 |
|  | S3 | 152.29 ±16.665 | 291.61 ±21.456 | 398.23 ±40.750 | 70.07 ±7.809 | 81.73 ±6.924 | 82.19 ±11.193 | 82.23 ±12.287 | 209.88 ±20.643 | 316.04 ±38.743 |
| PI 80831 | S1 | 34.44 ±3.224 | 125.13 ±19.150 | 465.64 ±64.862 | 13.07 ±2.169 | 26.46 ±5.553 | 72.98 ±7.450 | 21.36 ±3.129 | 98.67 ±17.386 | 398.85 ±63.330 |
|  | S2 | 147.16 ±20.161 | 306.19 ±41.438 | 779.37 ±35.609 | 48.09 ±13.082 | 60.27 ±8.330 | 102.54 ±14.586 | 103.47 ±17.694 | 245.92 ±39.476 | 676.82 ±37.134 |
|  | S3 | 246.57 ±14.062 | 322.54 ±27.929 | 559.50 ±43.416 | 107.34 ±5.283 | 111.57 ±13.125 | 153.20 ±12.519 | 139.24 ±13.042 | 210.97 ±22.288 | 406.31 ±40.598 |

**Table 4.** Shows the mean and standard error of the root and shoot dry biomass in three vegetative stages for each genotype and across three environments.

| Genotype | Env | Mean of Root Dry Mass (g) |  |  | Mean of Shoot Dry Mass (g) |  |  |
| --- | --- | --- | --- | --- | --- | --- | --- |
|  |  | V1 | V3 | V5 | V1 | V3 | V5 |
| CLOJ095-4-6 | S1 | 0.127 ±0.0064 | 0.371 ±0.0522 | 0.878 ±0.1152 | 0.524 ±0.0626 | 1.413 ±0.0766 | 3.700 ±0.4093 |
|  | S2 | 0.205 ±0.0149 | 0.458 ±0.0473 | 1.367 ±0.0972 | 0.724 ±0.0534 | 1.625 ±0.1398 | 4.867 ±0.2672 |
|  | S3 | 0.192 ±0.0078 | 0.354 ±0.0178 | 0.525 ±0.0422 | 0.345 ±0.0203 | 0.609 ±0.0506 | 1.439 ±0.0822 |
| PI 437462A | S1 | 0.083 ±0.0122 | 0.150 ±0.0500 | 0.600 ±0.0463 | 0.372 ±0.0415 | 1.238 ±0.2129 | 3.925 ±0.4769 |
|  | S2 | 0.143 ±0.0108 | 0.283 ±0.0477 | 0.978 ±0.1077 | 0.694 ±0.0415 | 1.411 ±0.1317 | 4.056 ±0.4628 |
|  | S3 | 0.165 ±0.0090 | 0.373 ±0.0232 | 0.516 ±0.0410 | 0.282 ±0.0206 | 0.525 ±0.0534 | 1.082 ±0.0890 |
| PI 438103 | S1 | 0.102 ±0.0140 | 0.225 ±0.0313 | 0.850 ±0.1014 | 0.347 ±0.0542 | 1.000 ±0.2204 | 4.280 ±0.6294 |
|  | S2 | 0.193 ±0.0136 | 0.367 ±0.0289 | 1.220 ±0.0512 | 0.717 ±0.0371 | 1.750 ±0.1003 | 5.430 ±0.2785 |
|  | S3 | 0.227 ±0.0214 | 0.359 ±0.0245 | 0.609 ±0.0423 | 0.361 ±0.0341 | 0.550 ±0.0597 | 1.327 ±0.0909 |
| PI 4381033B | S1 | 0.148 ±0.0204 | 0.501 ±0.1442 | 1.243 ±0.1494 | 0.492 ±0.0661 | 1.950 ±0.1793 | 6.014 ±0.4896 |
|  | S2 | 0.201 ±0.0230 | 0.724 ±0.0752 | 1.700 ±0.0928 | 0.870 ±0.0790 | 2.650 ±0.2070 | 6.225 ±0.3736 |
|  | S3 | 0.277 ±0.0139 | 0.483 ±0.0479 | 0.796 ±0.0684 | 0.462 ±0.0390 | 0.756 ±0.0705 | 1.766 ±0.2041 |
| PI 471899 | S1 | 0.142 ±0.0096 | 0.359 ±0.0506 | 1.050 ±0.1389 | 0.560 ±0.0482 | 1.825 ±0.1373 | 4.550 ±0.4633 |
|  | S2 | 0.180 ±0.0115 | 0.400 ±0.0333 | 1.110 ±0.1080 | 0.775 ±0.0703 | 1.889 ±0.0920 | 4.150 ±0.3027 |
|  | S3 | 0.230 ±0.0223 | 0.309 ±0.0187 | 0.489 ±0.0278 | 0.408 ±0.0252 | 0.568 ±0.0520 | 1.266 ±0.0825 |
| PI 80831 | S1 | 0.133 ±0.0144 | 0.333 ±0.0373 | 1.000 ±0.1174 | 0.482 ±0.0418 | 1.520 ±0.2086 | 5.630 ±0.5604 |
|  | S2 | 0.203 ±0.0154 | 0.540 ±0.0723 | 1.470 ±0.1136 | 0.835 ±0.0796 | 2.144 ±0.2789 | 5.700 ±0.4636 |
|  | S3 | 0.312 ±0.0141 | 0.441 ±0.0209 | 0.743 ±0.0577 | 0.502 ±0.0170 | 0.698 ±0.0462 | 1.716 ±0.2299 |

**Table 5.**
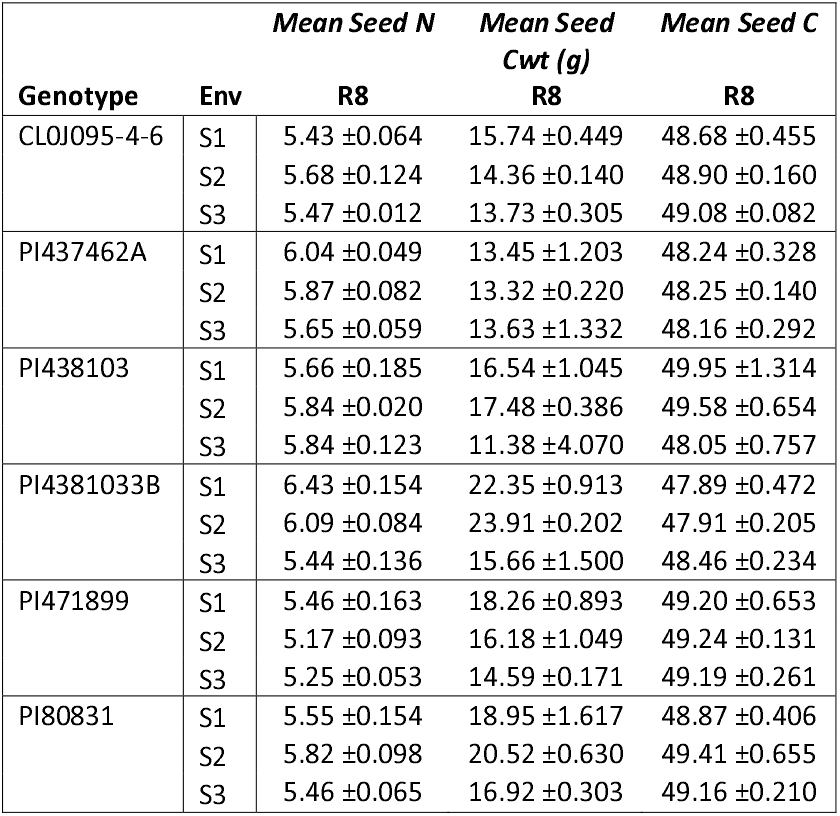
Shows the mean and standard error of Nitrogen (N), one hundred seed weight (Cwt), and Carbon (C) from mature seeds of six soybean accessions. Nitrogen and carbon are shown as a percent of the total seed weight, while the Cwt is shown in grams per 100 seeds.

When assessing all genotypes together, we can see that the taproot nodule counts are significantly influenced by the genotype and somewhat by the environment but not significantly influenced by the growth stage. However, the non-taproot growth zone was significant between each growth stage. This was confirmed by running a pairwise comparison of the growth stages of all genotypes and locations, comparing the count changes from stages V1 to V3, V3 to V5, and V1 to V5, all of which showed significance (p < 0.01). In contrast, the non-taproot nodule counts showed no significant changes between the growth stages in the taproot nodules (p > 0.1).

### 3.2 Differences in nodule count versus size lead to total area increases and differences in nodulation

The six genotypes evaluated across the three vegetative growth stages in the three environments resulted in similar observed trends in total nodule counts with differing variations each year. While the total nodule count increased with each advancement in the growth stage for all genotypes, not all genotypes increased nodule counts at the same rate. PI 438103 had the greatest increase of nodules between V1 to V3 with a rate of gain of 2.1x on average, while PI 4381033B had the least rate of gain at 1.5x between growth stages (Figure 2A). Cl0J095-4-6 had the greatest increase in noodle count from the V3 to V5 stage relative to the other genotypes, with a rate of gain of 2.3x. Genotypes PI 438133B and PI 471899 also showed a substantial increase in nodules from the V3 to V5 stage with average rates of gain at 2.2x, compared to the nodule count rate of gain in other genotypes. PI 473462A had the slowest rate of gain with 1.6x from V3 to V5. The total nodule count for the six genotypes at the V1 stage was similar (28-55 total nodules), while more apparent and discernable groups were established by V5, wherein, the six genotypes diverged into three groups, with two genotypes in the lower nodule count group (~88 nodules) and three genotypes in the middle group (~ 117 nodules). This third group consisted of PI 438133B (~166 nodules, Figure 2A). In the non-taproot nodules, there were two groups of nodule counts in the V1 stage [~21 nodules and ~37 nodules; Table 7(V1)], while non-taproot nodule counts of the genotypes shifted toward less discernable groups by V5 [Table 7(V5)].

**Figure 2:**
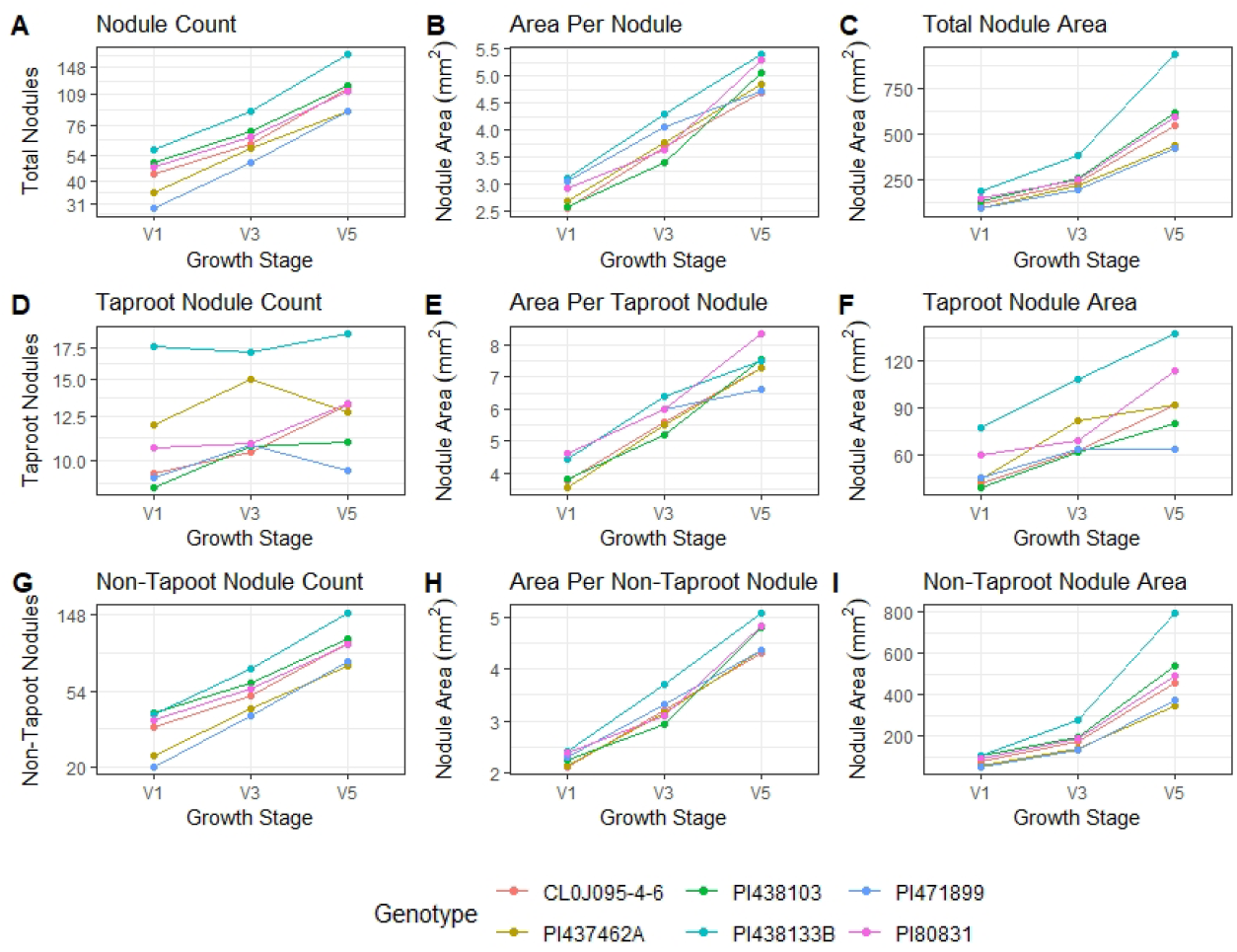
Average nodule count in the entire root, taproot, and non-taproot for six soybean accessions evaluated at three vegetative growth stages in three environments. All graphs show results for each genotype on a per plant basis at each growth stage. The number of replications per environment was 10-12. (A) total average nodule counts, (B) average individual nodule size (mm^2^), (C) total nodule volume, (D) total average taproot nodule counts, (E) average nodule size (mm^2^) of the nodules on the taproots, (F) total taproot nodule volume, (G) total average nodule counts of the non-taproot nodules, (H) average individual nodule size (mm^2^) of the non-taproot nodules, (I) average nodule size (mm^2^) of the nodules on the non-taproots.

When comparing the taproot nodule counts across each growth stage, it was observed that each genotype did not have consistent increases at each V stage. However, there were different growth patterns between the genotypes (Table 6). Two genotypes, PI 437462A and PI 471899, showed an increase in taproot nodulation from V1 to V3 and a significant drop in counts from V3 to V5 (Figure 2D). PI 80831 showed a minimal rise from V1 to V3, then increased by V5. In contrast, PI 438103 had increased nodule count from V1 to V3 and remained stable from V3 to V5. PI 438133B maintained nearly constant taproot nodule counts, while CL0J095-4-6 showed a near-constant increase in taproot nodule counts between each growth stage. Through changes and even some decreases in some genotype’s taproot nodule counts (Figure 2D), the individual taproot nodule volume compensated and continued to increase as the plant growth stages continued developing (Figure 2E), resulting in the total taproot nodule volume in most of the genotypes to increase (Figure 2F).

**Table 6.**
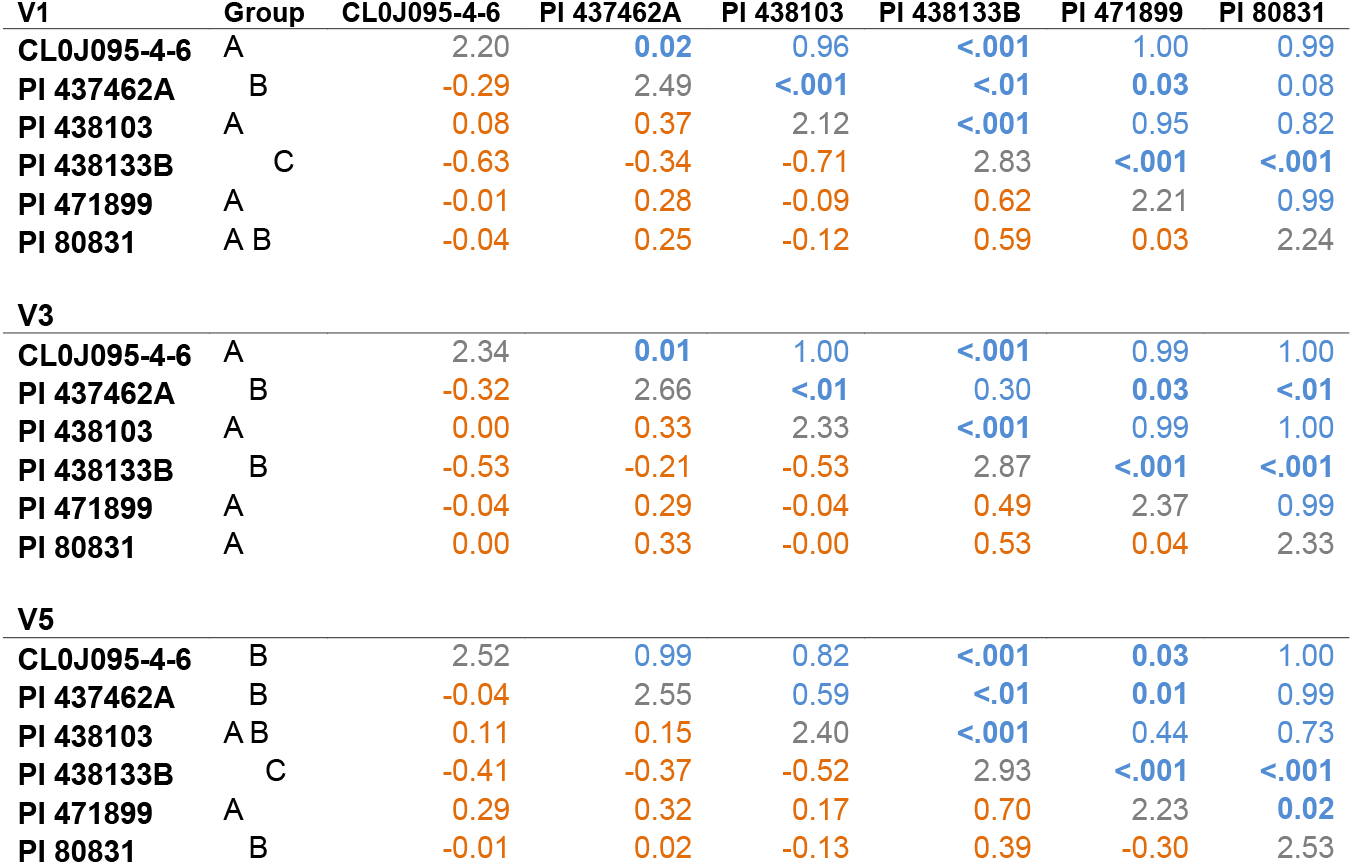
Pairwise comparisons of each soybean genotype taproot nodules (log-transformed) per growth stage showing Tukey adjusted p-values in the upper triangles (blue). The comparisons with significant differences at the 0.05 level are bolded. The table diagonals represent the mean estimates (gray). The lower triangles show the estimate comparisons (orange) calculated using earlier minus later comparisons. Group shows the groups of means that are not shown to be different at α = 0.05.

| V1 | Group | CL0J095-4-6 | PI 437462A | PI 438103 | PI 438133B | PI 471899 | PI 80831 |
| --- | --- | --- | --- | --- | --- | --- | --- |
| CL0J095-4-6 | A | 2.20 | **0.02** | 0.96 | **<.001** | 1.00 | 0.99 |
| PI 437462A | B | -0.29 | 2.49 | **<.001** | **<.01** | **0.03** | 0.08 |
| PI 438103 | A | 0.08 | 0.37 | 2.12 | **<.001** | 0.95 | 0.82 |
| PI 438133B | C | -0.63 | -0.34 | -0.71 | 2.83 | **<.001** | **<.001** |
| PI 471899 | A | -0.01 | 0.28 | -0.09 | 0.62 | 2.21 | 0.99 |
| PI 80831 | A B | -0.04 | 0.25 | -0.12 | 0.59 | 0.03 | 2.24 |
| V3 |  |  |  |  |  |  |  |
| CL0J095-4-6 | A | 2.34 | **0.01** | 1.00 | **<.001** | 0.99 | 1.00 |
| PI 437462A | B | -0.32 | 2.66 | **<.01** | 0.30 | **0.03** | **<.01** |
| PI 438103 | A | 0.00 | 0.33 | 2.33 | **<.001** | 0.99 | 1.00 |
| PI 438133B | B | -0.53 | -0.21 | -0.53 | 2.87 | **<.001** | **<.001** |
| PI 471899 | A | -0.04 | 0.29 | -0.04 | 0.49 | 2.37 | 0.99 |
| PI 80831 | A | 0.00 | 0.33 | -0.00 | 0.53 | 0.04 | 2.33 |
| V5 |  |  |  |  |  |  |  |
| CL0J095-4-6 | B | 2.52 | 0.99 | 0.82 | **<.001** | **0.03** | 1.00 |
| PI 437462A | B | -0.04 | 2.55 | 0.59 | **<.01** | **0.01** | 0.99 |
| PI 438103 | A B | 0.11 | 0.15 | 2.40 | **<.001** | 0.44 | 0.73 |
| PI 438133B | C | -0.41 | -0.37 | -0.52 | 2.93 | **<.001** | **<.001** |
| PI 471899 | A | 0.29 | 0.32 | 0.17 | 0.70 | 2.23 | **0.02** |
| PI 80831 | B | -0.01 | 0.02 | -0.13 | 0.39 | -0.30 | 2.53 |

Environment was significant for nodule count on non-taproots and taproots (see section 3.1); however, the main environmental effect was not significant for individual nodule size in the non-taproot zone (p = 0.11) and not significant for taproot individual nodule area (p = 0.13) (Supplemental Table S1). For total nodule area, environment was significant for both taproot (p = 0.01) and non-taproot (p = 1.02 × 10^−3^) growth zones (Supplemental Table S2). Taproot and non-taproot nodules varied in quantity between environments, with the lowest values consistently in S1 (Figure 1).

When we examined the effect of growth stage related to nodule count, this interaction was significant in the non-taproot nodules (p = 2.2 × 10^−16^) but not in the taproot nodules (p = 0.71). However, concerning the growth stage, we see that it is again highly influential on the average individual nodule size in the non-taproot growth zone (p = 1.68 × 10^−4^), and the individual nodule area was significant (p = 0.02) in the taproot zone as well. Significance of the growth stage effect was observed in both taproot (p = 0.05) and non-taproot zones (p = 3.21 × 10^−6^) for the average total nodule area (Figure S2).

Broadly speaking, different growth patterns between nodule taproot and non-taproot nodule quantities were observed (Tables 6, 7). Even though there is variation within genotypes, there is relatively low variation between genotypes in most environments (Figure 2). With some exceptions, Genotype PI 438133B consistently has a higher total nodule area than the other genotypes in all three growth stages (Figure 2 C F and I). This genotype also has some of the largest within genotype variations. PI 437462A had an exceedingly high nodule count at the V3 stage relative to all other genotypes and stages in S2 (Figure 3 D). PI 471899 was consistently lower for total nodule count than the rest of the genotypes, but over two years, it also showed an increase from V1 to V3 and then a decrease in nodule counts from V3 to V5 (Figure 3 D).

**Table 7.**
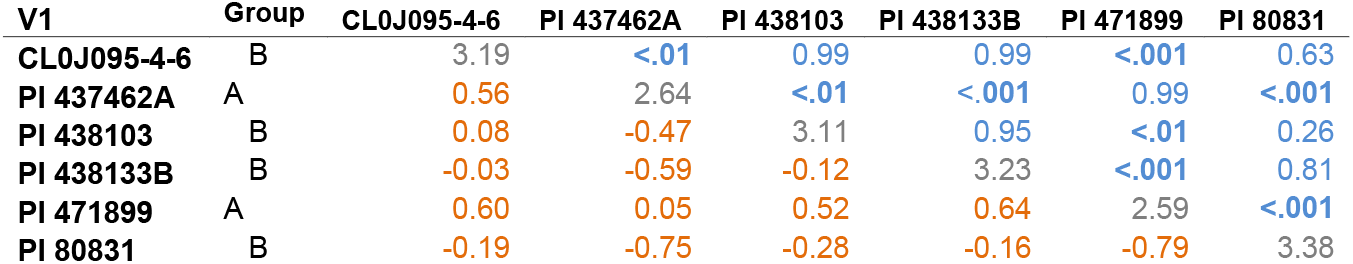
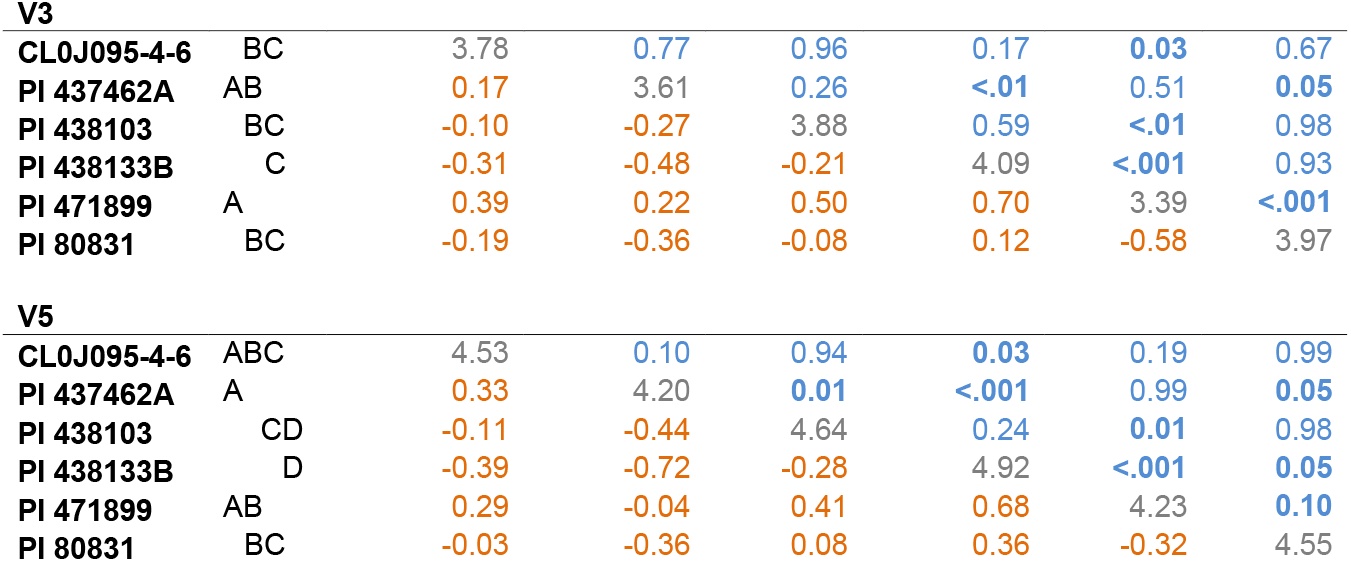
Pairwise comparisons of each soybean genotype non-taproot nodules (log-transformed) per growth stage showing Tukey adjusted p-values in the upper triangles (blue). The comparisons with significant differences at the 0.05 level are bolded. The table diagonals represent the mean estimates (gray). The lower triangles show the estimate comparisons (orange) calculated using earlier minus later comparisons. Group shows the groups of means that are not shown to be different at α = 0.05.

| V1 | Group | CL0J095-4-6 | PI 437462A | PI 438103 | PI 438133B | PI 471899 | PI 80831 |
| --- | --- | --- | --- | --- | --- | --- | --- |
| CL0J095-4-6 | B | 3.19 | **<.01** | 0.99 | 0.99 | **<.001** | 0.63 |
| PI 437462A | A | 0.56 | 2.64 | **<.01** | **<.001** | 0.99 | **<.001** |
| PI 438103 | B | 0.08 | -0.47 | 3.11 | 0.95 | **<.01** | 0.26 |
| PI 438133B | B | -0.03 | -0.59 | -0.12 | 3.23 | **<.001** | 0.81 |
| PI 471899 | A | 0.60 | 0.05 | 0.52 | 0.64 | 2.59 | **<.001** |
| PI 80831 | B | -0.19 | -0.75 | -0.28 | -0.16 | -0.79 | 3.38 |
| V3 |  |  |  |  |  |  |  |
| **CL0J095-4-6** | BC | 3.78 | 0.77 | 0.96 | 0.17 | 0.03 | 0.67 |
| **PI 437462A** | AB | 0.17 | 3.61 | 0.26 | <.01 | 0.51 | 0.05 |
| **PI 438103** | BC | -0.10 | -0.27 | 3.88 | 0.59 | <.01 | 0.98 |
| **PI 438133B** | C | -0.31 | -0.48 | -0.21 | 4.09 | <.001 | 0.93 |
| **PI 471899** | A | 0.39 | 0.22 | 0.50 | 0.70 | 3.39 | <.001 |
| **PI 80831** | BC | -0.19 | -0.36 | -0.08 | 0.12 | -0.58 | 3.97 |
| V5 |  |  |  |  |  |  |  |
| **CL0J095-4-6** | ABC | 4.53 | 0.10 | 0.94 | 0.03 | 0.19 | 0.99 |
| **PI 437462A** | A | 0.33 | 4.20 | 0.01 | <.001 | 0.99 | 0.05 |
| **PI 438103** | CD | -0.11 | -0.44 | 4.64 | 0.24 | 0.01 | 0.98 |
| **PI 438133B** | D | -0.39 | -0.72 | -0.28 | 4.92 | <.001 | 0.05 |
| **PI 471899** | AB | 0.29 | -0.04 | 0.41 | 0.68 | 4.23 | 0.10 |
| **PI 80831** | BC | -0.03 | -0.36 | 0.08 | 0.36 | -0.32 | 4.55 |

**Figure 3.**
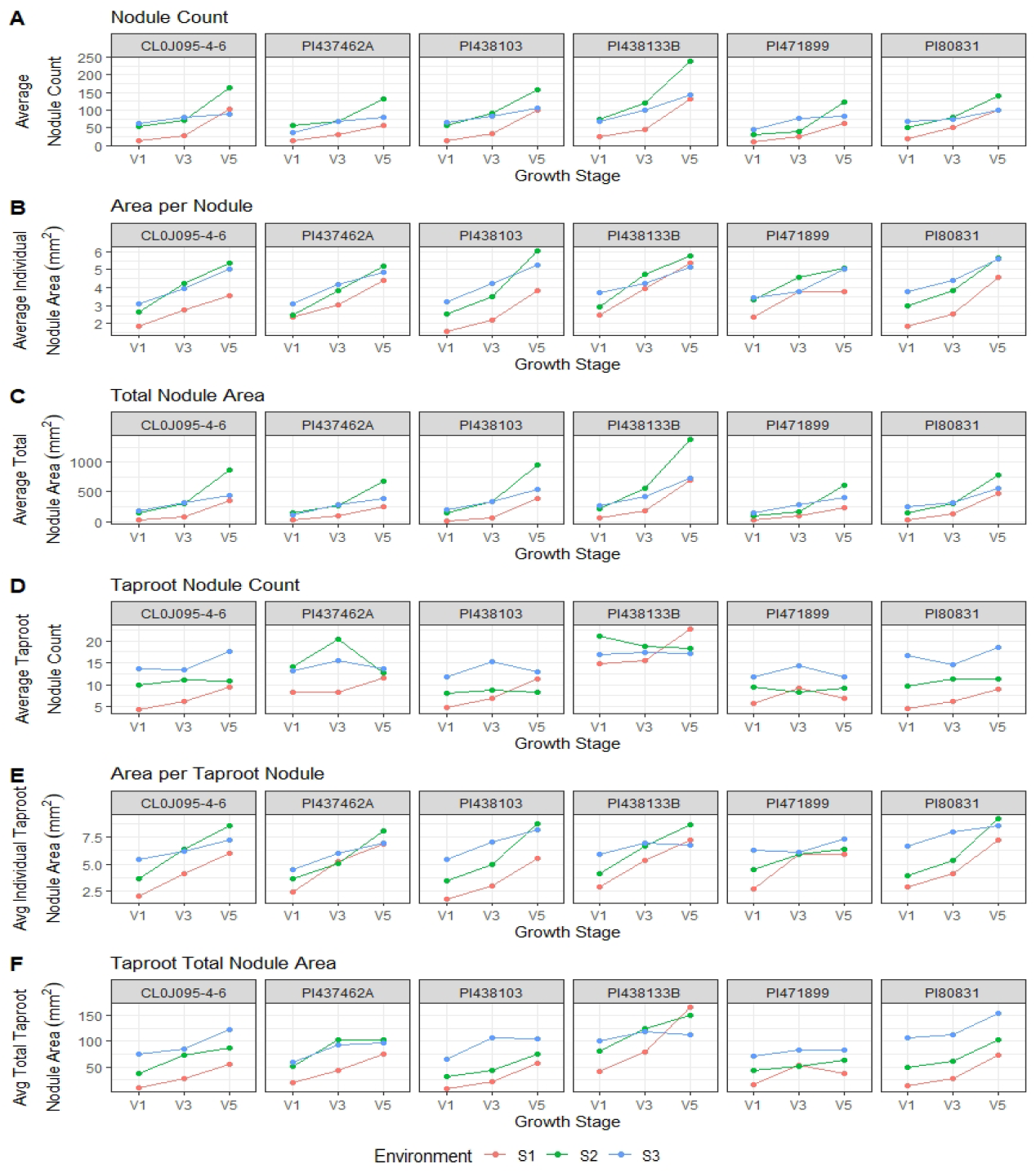
Nodule information on the entire root and taproot for six soybean accessions evaluated at three vegetative growth stages in three environments. (A) Average nodule count. (B) Average individual nodule area. (C) Average total nodule area. (D) Average taproot nodule count. (E) Average taproot individual nodule area. (F) Average total taproot nodule area.

Figure 3 shows each genotype’s nodule count, individual nodule area, and total nodule area by environment. While PI 438133B had the most taproot nodules, it was still comparable with the remaining genotypes in individual taproot nodule area (Figure 3 D and E), resulting in a higher total nodule area. We can also see the opposite effect where PI 471899 has fewer taproot nodules on average while remaining similar to the comparing genotypes in terms of total taproot nodule area due to an increased individual taproot nodule area. Cl0J095-4-6 and PI 80831 have greater variation in taproot nodule counts between environments resulting in more considerable variation in total taproot nodule area between environments.

On average, individual taproot nodule areas were larger at each growth stage than non-taproot individual nodule areas (Figure 3 B and E). We can see the average range for V1 non-taproot nodules ranges from 1.5-3.8 mm^2,^ while average taproot nodule areas range from 3.5-6.4 mm^2^. This range is further widened by V5, where we see the non-taproot nodule areas ranging from 3.5-6.1 mm^2^ and the taproot nodule areas ranging from 6.1-8.4 mm^2^ (Figure 3 B and E). While taproot growth is one of the first regions of the root to grow and set nodules, the nodules on the taproots increased in individual nodule area at a greater rate. On the other hand, nodules on the non-taproots are increasing in count and area.

### 3.3 Comparison of Nodulation Traits with Final Seed Nitrogen and Plant Biomass

There is a strong correlation between the total elemental nitrogen content as measured by the total percent N in the seed and taproot nodulation (Table 8). The average count of taproot nodules had the most consistent high correlation with N% in the final seed of the plants. For taproot nodules, no correlation was noted across all environments between individual nodule area and total seed N% (Figure 4). For non-taproot nodules, a low correlation was observed between individual nodule area and total seed N% at all early growth stages in S1 and S2 but not in S3 (data not presented).

**Table 8.** Select nodule traits and their correlation with the percent seed nitrogen. Pearson coefficient and associated p-value are presented. Data comes from six soybean genotypes grown in 10-12 replications in each of the three environments. S1 and S2 were two distinct environments in 2018, and S3 was grown in 2019.

| Trait | S1 & S2 |  | S1, S2, & S3 |  |
| --- | --- | --- | --- | --- |
|  | %N | P-value | %N | P-value |
| Taproot Nodulation |  |  |  |  |
| V1 | 0.42 | 0.17<sup>ns</sup> | -0.11 | 0.67<sup>ns</sup> |
| V3 | **0.52** | **0.08\*** | 0.16 | 0.53<sup>ns</sup> |
| V5 | **0.82** | **>0.001\*\*\*** | **0.50** | **0.04\*\*** |
| Taproot Nodule Count |  |  |  |  |
| V1 | 0.63 | 0.40<sup>ns</sup> | 0.28 | 0.26<sup>ns</sup> |
| V3 | **0.60** | **0.03\*\*** | 0.30 | 0.23<sup>ns</sup> |
| V5 | **0.82** | **0.04\*\*** | **0.44** | **0.07\*** |

\*\*\*, \*\*, \* Indicates significance at alpha equal to >0.001, 0.05 and 0.1 respectively. ns indicates not significant at alpha equal to 0.1.

**Figure 4.**
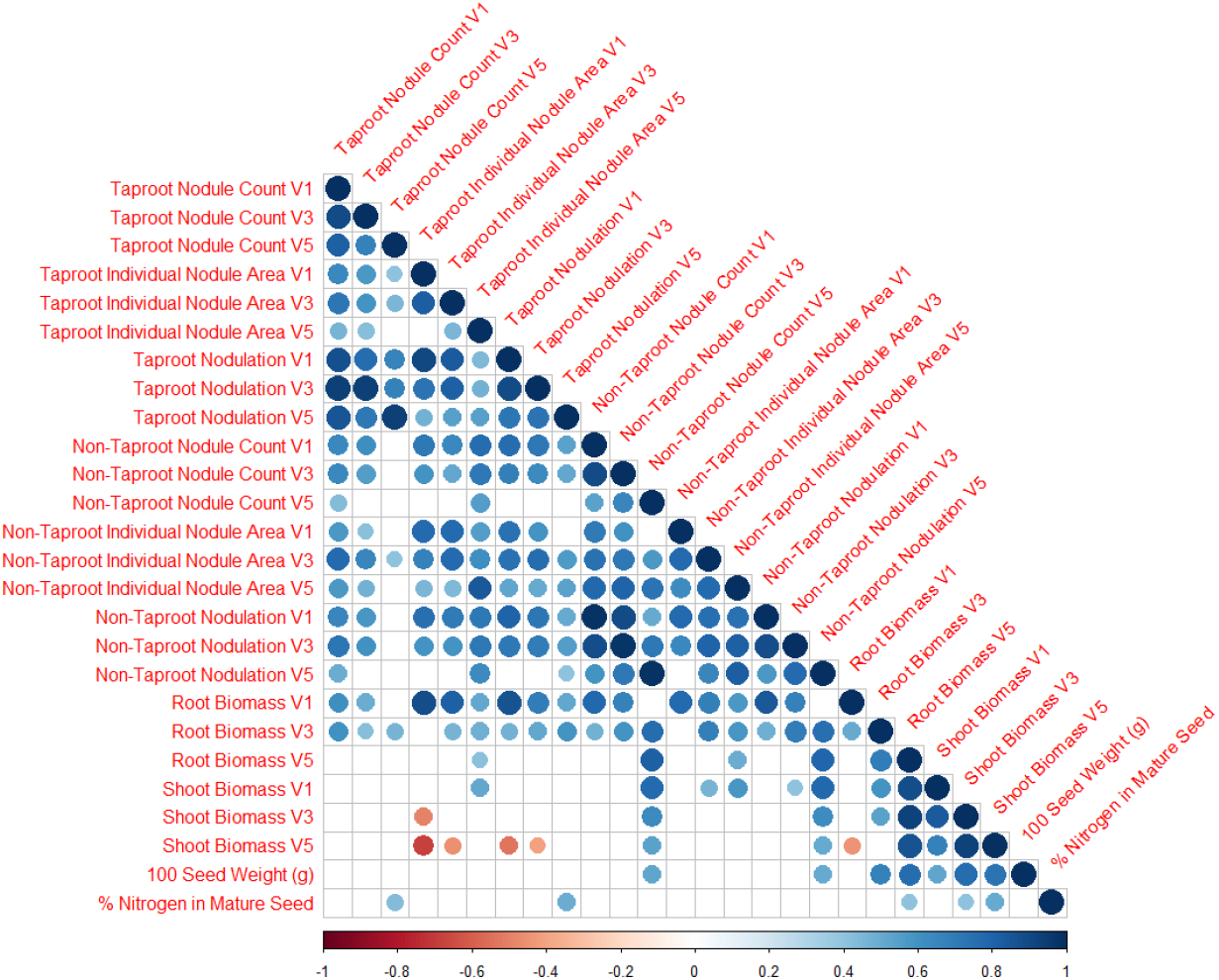
Spearman correlation of significant (α > 0.1) trait correlations for nodule traits. Data comes from six soybean genotypes grown in 10-12 replications in each of the three environments.

In 2018, on taproot, a significant correlation was noted at V3 and V5 for nodule area and count with seed N%; however, at the V1 growth stage, no correlation was observed between seed N% and nodule area or count (Table 8). When all three environment’s data were pooled, significant associations were only noted at the V5 growth stage between taproot nodule area and count with seed N%.

We parse out trait relationships into four categories: (a) nodule traits within taproots, (b) nodule traits across tap root and non-tap root, (c) nodule traits with root and shoot biomass, (d) shoot biomass and seed N% and 100 seed weight (gm).

For taproots, the correlation between individual nodule area and nodulation (i.e., total nodule area) diminished with increasing growth stage [at V1 (0.91, p < 0.001), at V3 (0.82, p < 0.001), and at V5 (0.55, p = 0.019)] (Table S3).

A significant correlation was observed between nodule count and nodulation, both on the taproot (0.93, p < 0.001) and non-taproot (0.98, p <0.001) at the V5 growth stage (Table S3), indicating that as the quantity of nodules increases, the total nodule area increases. We report that from V1 to V5 growth stages, taproot nodule count does not drastically change; however, an increase in individual taproot nodule area was observed. Together this may be causing the diminished or lack of correlation between nodule count and individual nodule area by V5. On the other hand, in the non-taproots, we noted a consistent correlation between nodule count and individual nodule area, as both traits showed a consistent relationship from V1 to V5 (Table S3).

Root and shoot biomass become increasingly correlated to each other with each growth stage (V1- 0.13, ns; V3- 0.53, p = 0.023; V5- 0.86, p < 0.001). There is a strong correlation between the root biomass in growth stage V5 and the non-taproot nodule count in growth stage V5 (0.81, p < 0.001), suggesting that with increased non-taproot growth, the count of non-taproot nodules increases as well. For taproots, the correlation between nodule count and root biomass diminished with increasing growth stage [at V1 (0.86, p < 0.001), at V3 (0.51, p < 0.029), and at V5 (0.16, p = 0.519)]. Whereas, for non-taproots the same correlation was strong through the growth stages [at V1 (0.84, p < 0.001), at V3 (0.69, p = 0.001), and at V5 (0.79, p < 0.001)]. Since in the early growth stages, the taproot is growing faster than the non-taproots, on average, nodules form and do not significantly change the count from V1 to V5. Whereas, in later growth stages, there is more non-taproot growth resulting in more non-taproot nodulation.

Shoot biomass and seed nitrogen content were also correlated across all environments at the V3 (0.41, p = 0.09) and V5 (0.50, p = 0.03) growth stages. Additionally, the Cwt of seeds were strongly correlated with root biomass at V5 (0.76, p < 0.001) and with shoot biomass at V3 (0.77, p < 0.001) and V5 (0.73, p < 0.001).

## 4. Discussion

### 4.1 Growth Stage and Environmental Impacts on Nodulation

In this work, we show evidence that unique genotypes have varying rates of nodule count in both the taproot and non-taproot growth zones (taproot p < 2.2 × 10-16, non-taproot p < 2.2 × 10-16) as each of these genotypes expresses a unique root system architecture. There is also varying response of nodule count in genotypes across environments (non-taproot p = 2.67 × 10-6, taproot p = 0.10) which also confirms prior understanding of environmental effects of nodule counts (Adekanmbi, 2019; Keyser & Cregan, 1987; Sadowsky et al., 1991; Sinclair et al., 1991; Wright, 1983). We also confirm reports of change in nodule count between growth stages of soybean root development, but these changes were only significant in the non-taproot growth zone in our data (taproot p = 0.74, non-taproot p = 2.72 × 10-9) (Brewin, 1991; Lawn & Brun, 1974). Additionally, the minimal evidence for interaction between genotype and growth stage (taproot p = 0.13, non-taproot p = 0.09) also shows that most genotypes have a consistent nodule growth trend across their growth stages. The significant interaction between genotype and environment (G x E) for both root growth zone nodule counts (taproot p = 5.72 × 10-11, non-taproot p = 3.45 × 10-6) shows that there is variation within genotypes across various environments. The observed lower nodule counts at environment S2 are likely a result of the environment having a loamy soil type with endogenously higher soil organic matter (3.5% OM at S1, 2.0% OM at S2 and S3) and likely, available nitrogen as soil available nitrogen has been shown to reduce symbiotic nitrogen fixation in soybean (Allos & Bartholomew, 1955, 1959).

When looking further into the relationships between seed weight and nitrogen in the seed, it was found that the two historically highest yielding lines, PI471899 and CL0J095-4-6, had some of the lowest amounts of nitrogen in the seed and per Cwt of seed (Figure S2 and S4). This supports previous findings in the literature as nitrogen is a primary component of seed protein, and traditionally, protein is negatively correlated with increased yield (Assefa et al., 2019; Mariotti et al., 2008). These lines could also be seen as the most nitrogen use efficient, resulting in the highest yield with some of the fewest nodules on both the taproot and non-taproots. PI 438103 was able to achieve moderate total nitrogen in the seed (Figure S4) while having a high level of taproot zone nodules and lower non-taproot zone nodules (Figure 2D, 2G), suggesting that it may have a higher conversion efficiency of nitrogen from the taproot for end seed nitrogen. However, one genotype, PI 438133B, showed an interesting relationship as it had the highest nodule counts on both the taproot and non-taproot zones along with the highest average total nitrogen and seed Cwt and percent nitrogen in the seed while still having a moderate yield compared to the other lines of this study (Figure S2, S3, and S4). This line may indicate that breeding for nitrogen and protein component traits such as nodule-related traits could improve the protein yield relationship and bring qualities lost through the founding effect of today’s modern germplasm back into the breeding pipeline.

The observed trend across genotypes was that the rate of nodule counts varies between different growth stages in the non-taproot growth zone and that nodule count within the taproot growth zone does not statistically differ between the early growth stages. These trends fall into line with the notion of host autoregulation of nodulation as controlling the quantity of nodule formation and regulation of where nodules are formed (Kosslak & Bohlool, 1984; Tanabata & Ohyam, 2014; Wang et al., 2014). However, our findings challenge the notion of many nodules being auto-regulated to form near the root crown as we see genetic diversity of nodule regulation across the growth zones (Reid et al., 2011). In modern production practices, many soybeans have inoculum applied to the seed at planting, leading to higher nodulation around the root crown (Bhuvaneswari et al., 1981; Calvert et al., 1984). As our study was dependent on soil available rhizobia, we found nodulation spread across the entire root structure and variable between growth zones.

### 4.2 Redefining Nodulation

For all genotypes, the average individual nodule area in mm^2^ increased at each growth stage on both taproot and non-taproots. However, apparent differences were observed in individual nodule area growth rates among genotypes. PI 438103 and PI 80831 had a higher growth rate from V3 to V5 than the other genotypes for individual nodule areas on taproot and non-taproots (Figure 2 E, H). Four of the six genotypes have higher total nodule area growth rates in the taproot growth zone from V3 to V5, and PI 437462A and PI 471899 have slower growth rates (Figure 2 F). These results suggest the impact of variation in the total count and individual nodule area on the total nodule area among these genotypes. It has been shown that nodule count and biological nitrogen fixation can have a strong relationship (Pitumpe Arachchige et al., 2020), suggesting that the ability to screen genotypes for nodule count can have a relational impact on nitrogen fixation. The size of nodules is also directly related to biological nitrogen fixation, as shown in Brun (Lawn & Brun, 1974).

Since the environment was most influential on nodule count and not on individual nodule area, it could be considered that one primary environmental influence could be the concentration of endogenous bacteria in each environment, as lower bacteria concentrations could lead to lower nodule counts from the requirement of bacterial root proximity for nodule initiation (Caetano-Anollés & Gresshoff, 1991; Ferguson et al., 2019). Future studies should also include quantifying local rhizobia populations with methods such as 16S rRNA analysis of the soil communities (Mayhood & Mirza, 2021; Shiro et al., 2013).

Logically, the taproot is fixed in a location, only growing deeper into the soil horizon where fewer nodules tend to form. Since nodules tend to grow most in the first 0-30 cm of soil (Hardarson et al., 1989; Voorhees et al., 1976), the increasing number of non-taproots with each growth stage in this zone will also lead to increased opportunity for nodule initiation. Future studies can benefit from studying the significance of the total nodule area being affected by the growth stage with additional growth stages and genotypes to show the biological relevance of nodule counts and the average individual nodule area with its limitations across time.

By using diverse accessions and leveraging image and ML-based nodule count, size, placement, and growth, we were able to generate insights that were previously not or less frequently attempted. Therefore, from a biological perspective, we propose that nodulation is defined by the total nodule area and is controlled as a function of both the nodule count and the individual area of the nodules. We also propose that different genotypes enable different rates of nodulation through these two separate controls of nodule growth and development, that is, through (a) nodule count and (b) individual nodule area resulting in the growth of total nodule area. This is particularly evident in the taproot growth zone and its relation to the nodulation among genotypes. For example, PI 80831 had average taproot nodule counts compared to the other genotypes, but it had the largest average individual taproot nodule size, which resulted in higher average nodulation than most genotypes. While we are proposing this paradigm shift, we are also cognizant of the value of assessing nodule health and its activity (Denison et al., 1991; Oono & Denison, 2010). Therefore, this too will need to be a future consideration in similar experiments and should also be the focus of non-invasive techniques, perhaps using digital phenotyping to ensure a higher throughput.

Logically, the success of nodulation to increase seed nitrogen content will be dependent on the shoot biomass as shoot photosynthates will be transferred to the roots and nodules to be utilized for nodule growth and development and returned as amino acids, ureides, and additional forms of bioavailable nitrogen (Carter & Tegeder, 2016; Kouchi & Higuchi, 1988; Thorpe et al., 1998).

### 4.3 Relationships Between Nodulation Traits, End Seed Nitrogen Content, and Biomass

As we define nodulation (i.e., total nodule area) as a function of the count and individual nodule area, it was interesting to note that in the early vegetative growth stages V1, V3, and V5, only nodulation and nodule count had significant correlation with seed N% while individual nodule area did not. Furthermore, in non-taproot, no correlation exists between the three nodule traits and seed N% (data not presented, see Figure 4). Together, these results with previous work (Carciochi et al., 2019; Hwang et al., 2014; Oono & Denison, 2010; Purcell & King, 1996) show a promising avenue to breed for higher total seed N% leveraging the nodule trait information reported in this paper. Due to the nature of this study and its focus on vegetative growth stage data collection, it is possible that this relationship of nodule traits in taproots and end-of-season seed N% extends to reproductive stages. From a practical viewpoint and existing imaging technologies, early growth stages are still more accessible to phenotype for RSA and nodule traits; therefore, the V stage results are valuable for practical applications and predictions. Additional work will need to be done to evaluate the optimum taproot nodulation – size, count, and placement, selecting cultivars with the best taproot nodulation, and possibly reducing non-taproot nodulation to minimize the nodule carbon sink could potentially maximize nitrogen use efficiency relative to seed nitrogen content. Future work should also include high throughput studies on the genetic frameworks that control taproot and non-taproot zone-specific RSA, nodulation, and relationships with carbon production (Montes et al., 2022), which could lead to a greater understanding and managing of soybean Nodulation Carbon to Nitrogen Production Efficiency (NCNPE), which we define as the maximum levels of nitrogen produced by nodulation in relation to carbon consumed by nodulation. Some work has been done showing that the total N content exported from harvested seed was greater than the N produced through biological nitrogen fixation, further leading to explore efforts on increasing carbon while evaluating nitrogen availability (Córdova et al., 2019). While this concept needs validation and testing, in this paper, we will use this concept to describe the potential efficiency of nodulation on legume plants. Additional work should also explore the intricate impacts of root system architecture and various seed inoculations on biological nitrogen fixation and seed nitrogen content (Carciochi et al., 2019). The nodulation-biomass correlations give further credence to the earlier assertion that genotypes have varying responses to nodulation. Some show an increase in the number of nodules, and others show an increase in individual nodule area to reach the amount of nodulation the plant requires.

### 4.4 Assessment of Nodulation Traits for Crop Breeding

One of the longer-term goals of this research is to utilize the results of this study for crop improvement using nodulation traits in the breeding pipeline. Therefore, we were interested in studying the stability of nodulation traits among genotypes across environments, nodule health, and genetic variation for taproot versus non-taproot nodulation and the carbon-nitrogen relationship in the context of nodulation.

We noted that soybean genotypes have differences in nodulation capabilities and variation in nodule presence and their development, i.e., individual nodule area and total nodule area. Since we were interested in a preliminary examination of nodule health, in a very small subset of plots at each environment (5-10 in each environment), nodules were physically cut with a sharp razor blade. We observed that even small nodules on the roots (< 3 mm^2^) had the typical red color of active nitrogen fixation (Virtanen & Laine, 1946); the brightest red colors were consistently observed in the nodules on the taproots, and the most prominent nodules on the non-taproots. While SNAP can count individual nodules and quantify their sizes and root growth zones, it is currently unable to differentiate which nodules are immature, actively fixing, or senescing nodules. Another follow-up research area will be the utilization of additional resources such as hyperspectral imaging to identify specific bands correlated to active fixation, similar to what researchers have shown to identify disease signatures in soybean (Nagasubramanian et al., 2018, 2019) or for plant component predictions (Chiozza et al., 2021). An AMMI type stability analysis was conducted to study the stability of genotypes for important nodule traits (Figure S5). CL0J096-4-6 was the most stable genotype across all three environments and in all three growth stages for nodulation in the non-taproot growth zone, individual nodule area, and total nodulation in the taproot growth zone (Figure S5). While trait stability is beneficial, it is not generally useful without associated performance traits such as seed yield (D. P. Singh et al., 2021). CL0J096-4-6 did not have the highest nodulation trait values but compared to the other five genotypes in this study, only PI 471899 is traditionally higher yielding than CL0J096-4-6 (Parmley, 2019). Interestingly, when evaluating the taproot nodulation, PI 471899 was highly stable for nodule count and individual area (Figure S5). As CL0J096-4-6 is the only genotype that has been selectively bred in the U.S., it can be hypothesized that plant breeders have unintentionally selected nodulation in tandem with yield and favorable above-ground traits. However, follow-up research is needed to confirm the role of indirect selection of nodulation traits when breeders select for seed yield. Future studies can be done using yield trial plot configuration to study nodulation traits with other agronomic production system traits.

There is a wide range of relationships still available within the soybean germplasm for nodulation traits. As shown earlier, there is potential for genotypes to have low taproot nodule counts with high non-taproot nodule counts (PI 437462A), while the opposite is also observed (PI 438103). Some genotypes appear to be more efficient at using the nodules to produce higher yields (CL0J095-4-6, PI 471899) (Parmley et al., 2019). In contrast, others (PI 438133B) may have higher nodule counts and moderate yield over lower historically yielding lines that still have moderate nodule counts (PI 80831, PI 438103) (Parmley et al., 2019), suggesting that selecting for nodule traits is still a viable option for breeders to implement in their programs. Due to the extreme value of nodulation, a corpus of researchers in inter-disciplinary teams needs to develop information and tools that enable breeding for nodulation traits to improve crop production. From a research viewpoint and scientific advances, nodulation traits will be ideal for Meta-GWAS studies as they will enable trait relationships leveraging published studies (J. M. Shook, Zhang, et al., 2021).

Previous literature has shown a correlation between nodulation and drought tolerance (W. Du et al., 2009; Fenta et al., 2014; Hwang et al., 2014; Kunert et al., 2016). With the continued concern of climate extremes becoming ever-present across the globe and these extremes creating more significant impacts on league growth and production, it is essential to explore if nodulation can contribute to or hinder plant resilience to these extremes. Further work should be done to evaluate if nodules are merely linked to drought-tolerant traits or if they are contributing to drought tolerance themselves. While nodules serve as a significant carbon sink (Parvin et al., 2020) and nitrogen source (Libault, 2014), further evaluations of root system architecture traits that provide additional environmental resilience could enable selective breeding strategies that leverage nitrogen, carbon, and water use efficiency in future cultivars.

With access to more automated nodule identification and quantification methods, researchers can now investigate the formation of nodules on roots in a time-series manner to comprehend the bacteria-plant symbiosis (Jubery et al., 2021). This is the first step towards understanding and subsequently optimizing nodule development and positions on roots to enable us to produce usable N efficiently and, further, seed protein. This is due to the result of a growing plant-based protein market (Sabaté & Soret, 2014; T. Zhang et al., 2021), and existing market demand for higher seed protein crops for feed purposes (Kim et al., 2019) and leghemoglobin, which can be derived from soybean nodules (Fraser et al., 2018). The potential to selectively breed for nodulation versus yield exists to meet new and unique commercial product demands on nodules. Understanding nodulation and its placement in roots across a plant’s growth and development cycle could potentially improve nitrogen use efficiency and protein resources in plants. While this study explored the early vegetative growth stages, soybeans continue to nodulate and fix nitrogen until and through the seed fill stage of development (Yuan et al., 2017). However, SNAP’s current high thruput abilities are hindered by its 2D image limitations. As imaging technology improves and data acquisition from 3D point clouds becomes higher throughput (Chiranjeevi, 2021) and more informative (Cox et al., 2022), additional information could be collected on the 3D root nodule root relationships in later growth stages.

Continued development of high throughput phenotyping capabilities of root traits and machine learning with explainability (Tryambak Gangopadhyay, Johnathon Shook, Asheesh K. Singh, Soumik Sarkar, 2019) will enable the acquisition of formerly unknown relationships, as shown in this work. Additional work is needed to parse out the genetic controls between nodule count and individual nodule size to unlock the selection potential for breeders to maximize nitrogen and carbon use efficiency in soybeans across diverse genotype by environment interactions and climates (Krause et al., 2022; J. Shook et al., 2021). Combining advances in phenotyping as presented here with improved methods of genomic prediction will further enable breeding advancements related to nodulation (de Azevedo Peixoto et al., 2017; J. M. Shook, Lourenco, et al., 2021). Work could also be done using a similar method to evaluate other pulse crops’ ability to nodulate differentially in taproot and non-taproot zones (Nagasubramanian et al., 2021), potentially using less hands-on annotation (Nagasubramanian et al., 2022). We also foresee and recommend that farmers utilize imaging-based tools such as SNAP, if they are packaged in a smartphone app, allowing them to study nodulation and further work on root health for system-wide analysis and farm-based applications.

## 5. CONCLUSIONS

This work proposes that nodulation is defined as total nodule area, a function of both nodule count and individual nodule area. We see phenotypic variation in the ability of some genotypes to regulate the overall taproot nodule area by increasing the nodule count and others by growing more in individual nodule area to achieve the same objective of nodulation. We found several significant trends in the relationships between soybean root system architecture and nodulation. Broadly speaking, nodule counts are fixed from a very early growth stage on the taproot of the soybean plant. In contrast, the non-taproot nodules continue to increase in the count and individual nodule area through diverse genotypic and environmentally dependent rates. We were also able to quantify the changes in nodule sizes and count relative to their location on taproots versus non-taproots finding that taproot nodules are increasing primarily in the individual area while non-taproot nodules are increasing in both area and count. Further, we observed differences in genotype rates of nodulation on both taproots and non-taproots. In early growth stages, taproot nodulation and taproot nodule count correlate with mature seed nitrogen content. Additionally, root and shoot biomass at V5 correlated to total seed nitrogen at maturity.

We noted genotypes that had a stable performance in nodulation traits across environments, while other genotypes were unstable. Taproot traits tended to be more stable than non-taproot traits across environments. As we selected six diverse genotypes and three environments for this initial exploration, it is clear that additional work is needed to explore the soybean collection with greater depth to more robustly classify the nodule growth patterns in soybeans and across additional environments to explore further the G x E interactions and plant responses to weather. Breeders could leverage these genetic and environment interactions to build more stable performing lines or find optimal nodulating lines for specific environments in prescriptive breeding. Assessing nodule health, quality, and the nodulation carbon to nitrogen production efficiency, or NCNPE, are traits to be still considered. Therefore, this too will need to be a future consideration in similar experiments and should also be the focus of non-invasive techniques, perhaps using digital phenotyping to ensure a higher throughput. Our findings provide avenues to utilize nodulation traits in crop improvement.

## Author Contributions

Conceptualization - C.C. with A.K.S.; Methodology - C.C., M.Z., A.K.S.; Experimentation – C.C. and M.Z.; Statistical analysis - C.C. with S.D. and A.K.S.; Writing original draft - C.C. with A.K.S; Review and Editing – all authors; Research funding acquisition - A.K.S.; Project management - A.K.S.

## Funding

This project was supported by the Iowa Soybean Research Center, Iowa Soybean Association, R.F. Baker Center for Plant Breeding, Plant Sciences Institute, Bayer Chair in Soybean Breeding, USDA/NIFA Hatch CRIS Project IOW04717, USDA/NIFA Hatch Project IOW03717, USDA-NIFA #2021-67021-35329, NSF CPS#1954556, NSF S&CC#1952045. C.N.C. was partially supported by the National Science Foundation under Grant No. DGE-1545453.

## Acknowledgments

The authors thank the many undergraduate and graduate students who helped dig the roots and assisted in imaging them and counting nodules. Special thanks to the Singh Lab Staff members, particularly Ms. Jennifer Hicks and Mr. Brian Scott, for their assistance throughout the project that enabled project completion.

## Conflicts of Interest

The authors declare no conflict of interest.

## Data and code sharing

All data and code will be shared through the Singh group GitHub https://github.com/SoylabSingh.

## Supplemental Materials

Supplemental figures are available in the “Nodulation Supp Figures” document showing a visual representation of the roots used along with graphs of the percent nitrogen, hundred weight, and grams of nitrogen per hundred weights of seed. Additionally, a supplemental figure is provided showing the AMMI analysis with the description of how it was conducted and corresponding references. Supplemental tables are available in the “Nodulation Supp Tables” document and show the analysis of variances for individual nodule area and nodulation along with a table of the discussed correlations.

## Abbreviations

The following abbreviations are used in this manuscript:

IPCA: Interaction Principal Component Axis
N: Nitrogen
NCNPE: Nodulation Carbon to Nitrogen Production Efficiency (NCNPE
SNAP: Soybean Nodule Acquisition Pipeline
RSA: Root System Architecture
V(#): Vegetative growth stage followed by the number of the stage

## Notes

### Competing Interest Statement

The authors have declared no competing interest.

