## Supplemental Figures for "Using Machine Learning Enabled Phenotyping To Characterize Nodulation In Three Early Vegetative Stages In Soybean"

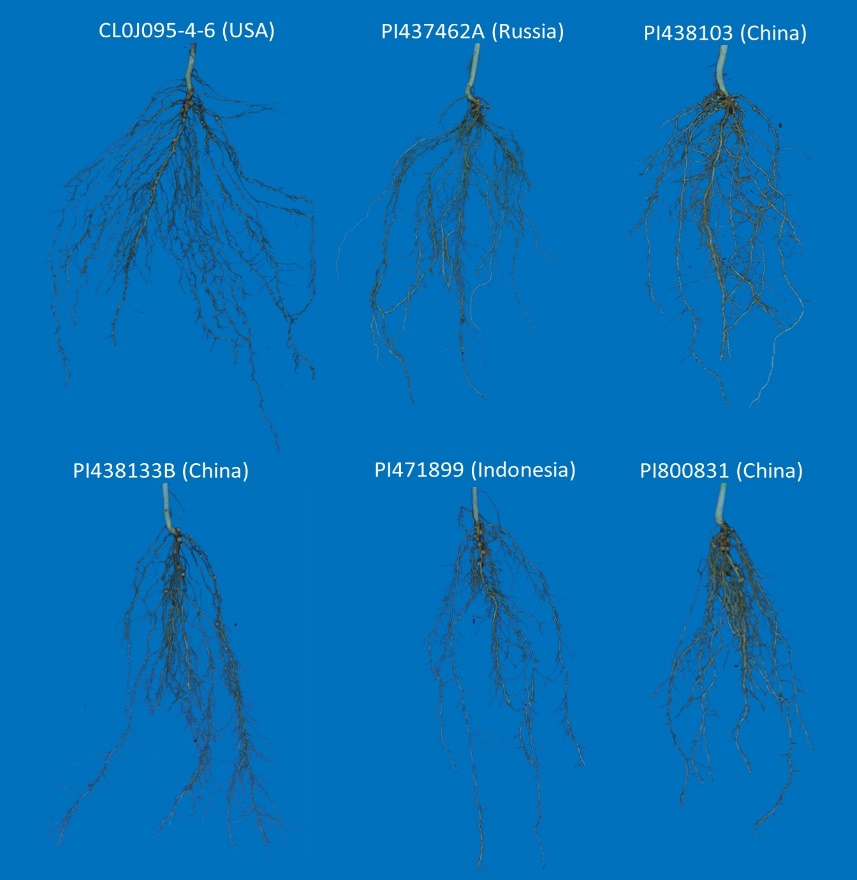

**Figure S1)** Representative images and countries of origin of the root phenotypes used to visually select the six genotypes evaluated in this study. Genotypes were selected to maximize genomic and phenomic dissimilarity while minimizing maturity and growth stage differences.

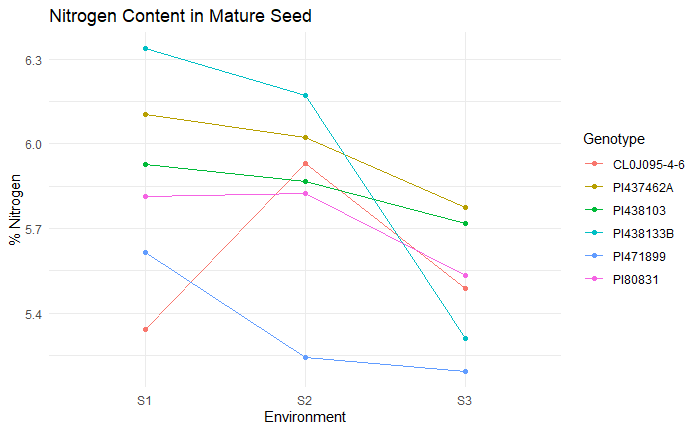
**Figure S2)** Average percent Nitrogen in mature seed (R8) of each genotype in each environment.

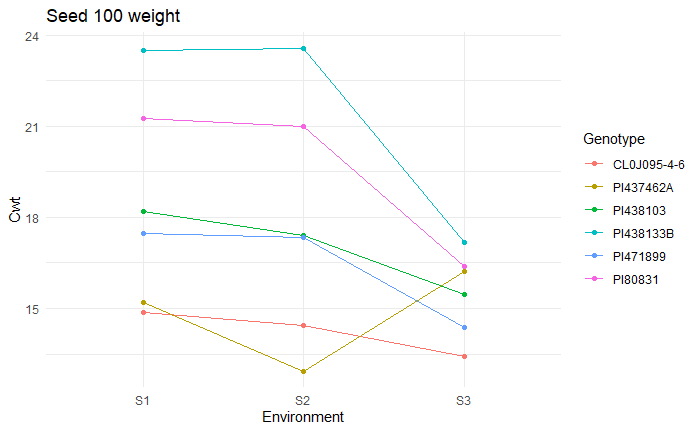
**Figure S3)** Average Seed Weight (g) of 100 mature seeds (R8) of each genotype in each environment.

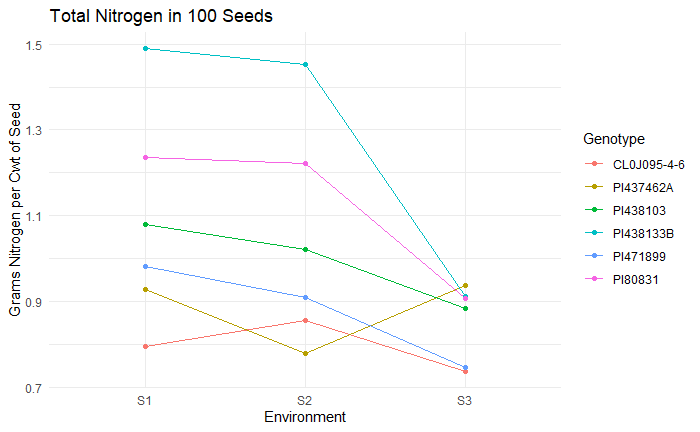
**Figure S4)** Average total Nitrogen in 100 mature seeds (R8) of each genotype in each environment as calculated by multiplying the percent Nitrogen from each line by the weight (g) of 100 seeds.

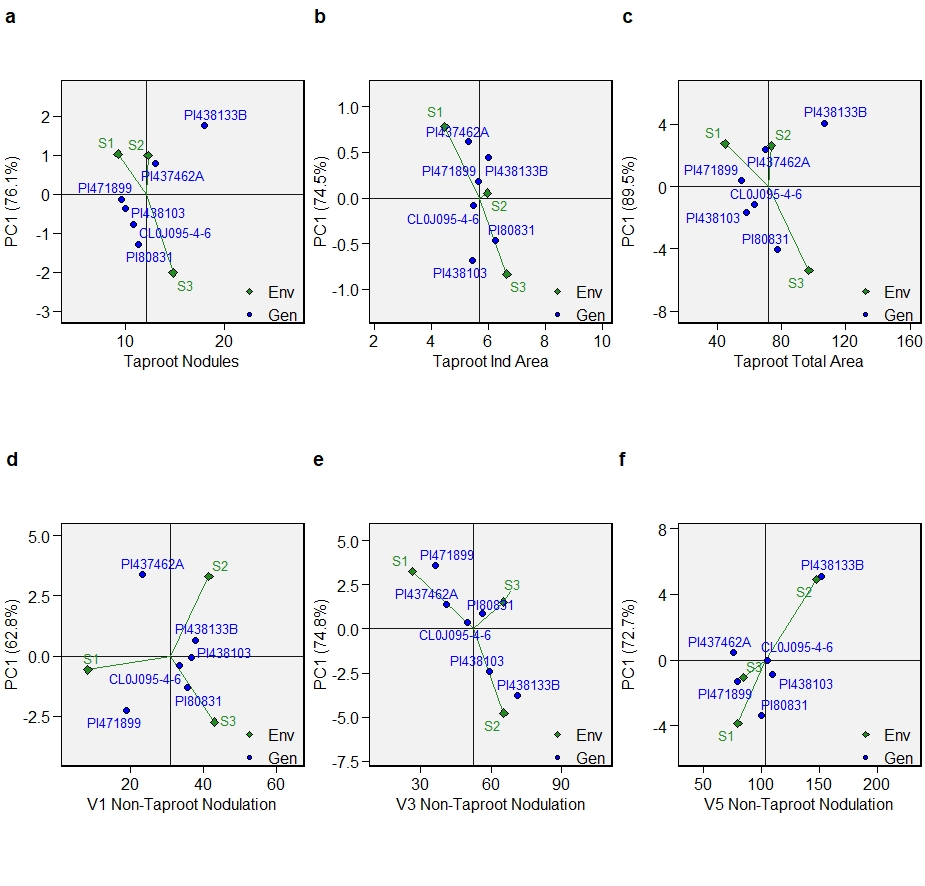

**Figure S5)** AMMI1 biplots for additive effects vs. IPCA1 in six genotypes across three environments. Panel A) Shows the taproot nodule count, B) The average individual taproot nodule area, and C) The taproot nodulation or average total taproot nodule area. Panels D, E, and F show the V1, V3, and V5 growth stages of non-taproot nodulation or taproot nodule area, respectively.

**Trait Stability Analysis**

AMMI analysis was conducted by adjusting the additive genetic effects and environmental effects through analysis of variance along with the adjustment of multiplicative effects from the G x E interactions through principal component analysis. The AMMI model used is as follows:

$Y_{ij}= \mu+ g_{i} + e_{j} + \sum_{k=1}^{p} \lambda_{k}a_{ik}t_{jk} + \rho_{ij} +\varepsilon_{ij}$	(S1)

Where *Y_ij_* is the nodule count of the *i^th^* genotype in the *j^th^* environment, *µ* is the overall mean, *g_i_* and *e_j_* are the fixed varietal effects and environmental deviations, respectively, *λ_k_* is a singular value of the *k^th^* axis in the interaction principal component analysis, *a_ik_* and *t_jk_* are genotype and environmental factors, respectively, of the singular eigenvectors associated with *λ_k_* from the interaction matrix. *P* is the number of principal components retained in the model, *ρ_ij_* is the residual *G×E* interaction if not all p IPCA are used, where *p ≤ min(g−1;e−1)*, and *ε^ij^* is the average independently assumed error *ε_ij_ ~N(0, σ^2^).*(Oliveira, Freitas, and Jesus 2014; Olivoto and Lúcio 2020)

Biplot graphs of the AMMI1 showing the first Interaction Principal Component Axis (IPCA1) versus additive effects from genotypes and environment (Figure S5). The S1 environment in 2018 consistently resulted in the lowest nodule counts in the taproot and non-taproot zones. While S2 and S3 gave the higher nodule counts in all three growth stages and both growth zones, we see a slight variation between S2 and S3 for producing more nodules between the growth stages, with them producing similar to or more non-taproot nodules in V1 and V3 but S2 producing far more in V5 (Figure S5 D, E, and F). However, S3 consistently had more taproot nodules.

In AMMI1, stability is evaluated in the y-axis (Duarte and Vencovsky 1999), with the most stable varieties being near the origin of the PCA1 axis. We can see that PI 471899, PI 438103, and CL0J096-4-6 appear to have the most stable taproot nodule counts across the environments, while PI 80831, PI 437462A, and PI 438133B have the most variation between environments. When looking at the non-taproot nodules, we see that CL0J096-4-6 is the most stable genotype across all three environments and in all three growth stages, while the PIs have much more variation. CL0J096-4-6 is a line that has historically been selectively bred for stability across environments as an elite line which could lead to the robust stability we also see in nodulation in this line. PI 438103 and PI 80831 are also seen to be somewhat consistently stable in the non-taproot growth zones, while PI 471899 is the most stable for taproot nodule count, individual nodule area, and nodulation.
