## Supplemental Tables for "Using Machine Learning Enabled Phenotyping To Characterize Nodulation In Three Early Vegetative Stages In Soybean"

| **Table S1** Type III Analysis of Variance of Taproot and Non-Taproot individual nodule area with Kenward-Roger's method. | | | | | |
| --- | --- | --- | --- | --- | --- |
|  |  | *Taproot Ind. Area* | | *Non-Taproot Ind. Area* | |
| **Source of variation** | **dF** | **F Value** | **P value** | **F Value** | **P Value** |
| Genotype (G) | 5 | 7.84 | 4.61 x 10^-7^*** | 7.12 | 2.15 x 10^-6^*** |
| Environment (E) | 2 | 1.98 | 0.13 | 2.52 | 0.08* |
| Growth Stage (T) | 2 | 3.73 | 0.02** | 7.37 | 6.25 x 10^-4^*** |
| G x E | 10 | 4.73 | 2.03 x 10^-6^*** | 3.66 | 1.10 x 10^-4^*** |
| G x T | 10 | 3.09 | 8.21 x 10^-4^*** | 2.48 | 6.79 x 10^-3^** |
| E x T | 4 | 0.34 | 0.85 | 0.08 | 0.99 |
| G x E x T | 20 | 1.17 | 0.27 | 2.10 | 3.87 x 10 ^-3^** |

Significance at *** < 0.001, ** 0.05, *0.1

| **Table S2** Type III Analysis of Variance of Taproot and Non-Taproot nodulation (total nodule area) with Kenward-Roger's method. | | | | | |
| --- | --- | --- | --- | --- | --- |
|  |  | *Taproot Nodulation* | | *Non-Taproot Nodulation* | |
| **Source of variation** | **dF** | **F Value** | **P value** | **F Value** | **P Value** |
| Genotype (G) | 5 | 35.21 | < 2.2 x 10^-16^*** | 24.67 | < 2.2 x 10^-16^*** |
| Environment (E) | 2 | 4.61 | 0.01** | 6.89 | 1.02 x 10^-3^** |
| Growth Stage (T) | 2 | 3.08 | 0.05* | 12.65 | 3.21 x 10^-6^*** |
| G x E | 10 | 9.86 | 8.18 x 10^-15^*** | 3.50 | 1.97 x 10^-4^*** |
| G x T | 10 | 3.57 | 1.54 x 10^-4^*** | 2.14 | 0.02** |
| E x T | 4 | 0.47 | 0.75 | 0.85 | 0.48 |
| G x E x T | 20 | 1.36 | 0.13 | 1.56 | 0.06* |

Significance at *** < 0.001, ** 0.05, *0.1

**Table S3** Correlations between Nodule Count, Individual Nodule Area, and Total Nodule Area for Taproot and Non-Taproot Growth Zones. Correlations are shown where p > 0.001, p > 0.05, p > 0.1, and p < 0.1 are represented by ***, **, *, and ns respectively.

|  | **V1 Ind Taproot Nodule Area** | **V3 Ind Taproot Nodule Area** | **V5 Ind Taproot Nodule Area** |
| --- | --- | --- | --- |
| **V1 Total Taproot Nodule Area** | 0.91*** | 0.84*** | 0.45* |
| **V3 Total Taproot Nodule Area** | 0.73*** | 0.82*** | 0.47** |
| **V5 Total Taproot Nodule Area** | 0.47** | 0.55** | 0.55** |
|  | **V1 Taproot Nodule Count** | **V3 Taproot Nodule Count** | **V5 Taproot Nodule Count** |
| **V1 Total Taproot Nodule Area** | 0.88*** | 0.77*** | 0.67** |
| **V3 Total Taproot Nodule Area** | 0.93*** | 0.94*** | 0.68** |
| **V5 Total Taproot Nodule Area** | 0.86*** | 0.72*** | 0.93*** |
|  | **V1 Taproot Nodule Count** | **V3 Taproot Nodule Count** | **V5 Taproot Nodule Count** |
| **V1 Ind Taproot Nodule Area** | 0.64** | 0.57** | 0.4* |
| **V3 Ind Taproot Nodule Area** | 0.72*** | 0.6** | 0.45* |
| **V5 Ind Taproot Nodule Area** | 0.47** | 0.43* | 0.24^ns^ |
|  | **V1 Ind Non-Taproot Nodule Area** | **V3 Ind Non-Taproot Nodule Area** | **V5 Ind Non-Taproot Nodule Area** |
| **V1 Total Non-Taproot Nodule Area** | 0.77*** | 0.75*** | 0.75*** |
| **V3 Total Non-Taproot Nodule Area** | 0.65** | 0.81*** | 0.82*** |
| **V5 Total Non-Taproot Nodule Area** | 0.35^ns^ | 0.65** | 0.83*** |
|  | **V1 Non-Taproot Nodule Count** | **V3 Non-Taproot Nodule Count** | **V5 Non-Taproot Nodule Count** |
| **V1 Total Non-Taproot Nodule Area** | 0.99*** | 0.89*** | 0.51** |
| **V3 Total Non-Taproot Nodule Area** | 0.88*** | 0.98*** | 0.72*** |
| **V5 Total Non-Taproot Nodule Area** | 0.60** | 0.72*** | 0.98*** |
|  | **V1 Non-Taproot Nodule Count** | **V3 Non-Taproot Nodule Count** | **V5 Non-Taproot Nodule Count** |
| **V1 Ind Non-Taproot Nodule Area** | 0.70*** | 0.61** | 0.25^ns^ |
| **V3 Ind Non-Taproot Nodule Area** | 0.75*** | 0.71*** | 0.57** |
| **V5 Ind Non-Taproot Nodule Area** | 0.79*** | 0.81*** | 0.73*** |
